# Multi-Enzyme Data-Independent Acquisition Mass Spectrometry for Proteoform-Aware Single-shot Proteomics

**DOI:** 10.64898/2026.11.06.764873

**Authors:** Moritz Heusel, Lotta Happonen, Christofer Karlsson, Emma Åhrman, Johan Malmström

## Abstract

Biological regulation does not operate at the level of the protein but across ensembles of protein variants shaped by alternative splicing, post-translational modification and endogenous proteolytic processing. Yet most conventional bottom-up discovery proteomics workflows remain optimized for a canonical protein census, leaving much of the sequence-level variation invisible. Multi-enzyme digestion has been recognized as an effective strategy to expand sequence coverage and proteoform sensitivity, but existing implementations rely on resource-heavy per-enzyme liquid chromatography coupled to mass spectrometry (LC-MS) analysis, dedicated measurements for spectral library construction, and semi-manual reconciliation across multiple software tools, restricting the use to specialized laboratories and limited sample cohorts. Here we introduce mE-DIA, a <u>m</u>ulti-<u>e</u>nzyme, proteoform-resolved <u>d</u>ata independent <u>a</u>cquisition (DIA)-MS workflow that multiplexes complementary protease digested samples into a single DIA injection and processes them through a library-free pipeline within a single analysis engine, without supplemental data dependent acquisition (DDA)-MS measurements. The mE-DIA Pipeline graphical user interface orchestrates the analysis, and a dedicated proteoform inference algorithm and the interactive mE-DIA Peptide Mapper viewer make the sequence-level evidence explorable. Comparing the workflow to prevailing DDA-library-based multi-enzyme DIA on the same raw data and within shared search space showed increased identified protein groups by 30 % and protein groups quantified at ≤ 20 % CV by 47 %. A three-enzyme multiplex of the human pathogen *Streptococcus pyogenes* resolved 990 protein-backbone variants among 309 canonical protein entries, 1.8-fold more than the most productive single-enzyme analysis, including an H69 C-terminal proteoform of the ribosomal protein uS19 that the trypsin-only workflow did not resolve. By consolidating multi-enzyme acquisition, analysis and proteoform interpretation into one accessible toolset, mE-DIA brings sub-gene-resolved evidence within reach of routine single-shot discovery DIA proteomics.

## Introduction

Biological systems are regulated at a level far more granular than individual proteins produced from a given genomic locus but rather ensembles of protein variants, termed ‘proteoforms’, per each gene of origin. The landscape of proteoforms is shaped by alternative splicing, posttranslational modification, and endogenous proteolytic processing. Estimates suggest that the combinatorial diversity of proteoforms in a single human sample may approach or exceed one million distinct molecular species. Consequently, analytically meaningful characterization of biological regulation requires methods capable of resolving variation across this landscape rather than collapsing it into a single inferred protein identity (Aebersold et al., 2018). Liquid chromatography coupled to mass spectrometry (LC-MS)-based proteomics is, in principle, capable of dissecting such complexity through targeted, problem-specific method development targeting proteoform-signature peptides (Korchak et al., 2024; Lange et al., 2008; Lin et al., 2023; Varela and Schmidt, 2022) or backbones from middle-or top-down perspectives (Huang et al., 2024; Toby et al., 2016), yet conventional and most frequently employed discovery proteomics workflows remain limited with regards to the discovery of specific protein variants and corresponding regulatory events.

High-throughput parallel affinity-based assay platforms, alike MS-based discovery workflows, optimize proteome coverage at gene-level counts, obscuring that these technologies are orthogonal to LC-MS-based proteomics and enrich rather than replace the proteomics toolbox (Beimers et al., 2025; Katz et al., 2022; Kirsher et al., 2025; Sissala et al., 2025). While affinity-based platforms have demonstrated biological relevance through pQTL mapping (Eldjarn et al., 2023; Sun et al., 2023) and high throughput scalable to 10,000s of clinical specimens, LC-MS-based proteomics is evolving rapidly through innovations in liquid chromatography (Bache et al., 2018; Falkenby et al., 2014), mass spectrometric equipment (Guzman et al., 2024; Heil et al., 2023) and DIA-MS, (Derks et al., 2023; Gillet et al., 2012; Silva et al., 2006; Venable et al., 2004), and has over the last 15 years significantly increased coverage of the canonical proteome, optimizing for the number of genes for which evidence could be detected in the single-enzyme-digest bottom-up LC-MS data. However, both affinity and conventional bottom-up MS-based discovery approaches fail to capture proteomes at the level of functional proteoforms, reporting either epitope-level signals that conflate concentration with accessibility and cross-reactivity or averaging peptide signals towards canonical protein abundance estimates. Both workflows leave opportunities to measure proteomes at a more granular level of functionally relevant proteoforms, an information partially embedded in but not commonly leveraged from the bottom-up LC-MS proteomic data.

While comprehensive characterization of intact proteoforms is traditionally the domain of top- and middle-down proteomics, these approaches remain challenged by proteome-scale application due to precursor complexity, extensive charge-state overlap, and the dynamic range of biological samples (Huang et al., 2024; Toby et al., 2016). Conventional bottom-up workflows, by contrast, achieve scalability and analytical robustness by reducing proteins to peptides, thereby simplifying chemical complexity at the expense of preserving long-range molecular context and proteoform connectivity.

Here, we sought to unlock opportunities to extend scalable bottom-up discovery proteomics by the ability to recover biologically meaningful proteoform-resolved information.

Multi-enzyme proteomics has been recognized as an effective strategy to expand sequence coverage and increase sensitivity to splice variants, endogenous proteolysis, and post-translational modification-dependent peptide observability (Guo et al., 2014; Sinitcyn et al., 2023). However, conventional implementations typically require separate LC-MS measurements and semi-manual downstream analyses for each protease (Richards et al., 2022), or focus on peptide mapping in low-complexity samples (Morsa et al., 2019), each limiting applicability in large-scale discovery settings.

Over the last 10-15 years, data-independent acquisition has become the preferred mode of label-free MS proteomics data acquisition once software advances allowed its inherent all-ion sampling to translate into improved quantitation as well as improved proteome coverage (Demichev et al., 2018; Gillet et al., 2012; Röst et al., 2014; Venable et al., 2004), reducing dependence on chromatographic gradient length to capture sample complexity and instrument speed to scan individual peptide precursor ions (Bruderer et al., 2017; Kelstrup et al., 2018; Schebesta et al., 2025). Motivated by our work on multi-enzyme deep learning models for *de novo* sequencing (Gueto-Tettay et al., 2023), we hypothesized that DIA-MS could effectively record multiplexed multi-enzyme peptide maps that may subsequently be interpreted either through *de novo* sequencing, or, alternatively, through enzyme-specific, peptide-centric de-multiplexing analysis. The latter approach was since demonstrated by a recent effort combining enzyme-multiplex DIA with per-enzyme DDA-MS measurements and optimization of peptide fragmentation conditions to cumulatively enhance protein sequence coverage (Richards et al., 2022). That study established the feasibility of peptide-centric enzyme de-multiplexing, but did so from separate per-enzyme DDA-MS measurements, merging the resulting enzyme-specific spectral libraries for concurrent query, with semi-manual analysis across three computational tools (Richards et al., 2022).

The work reported here shares that acquisition concept but differs in its analysis premise by not requiring separate DDA measurements, experimental spectral libraries or reconciliation across computational tools, with enzyme de-multiplexing and proteoform inference performed in a single analysis framework. The mE-DIA workflow is a multi-enzyme extension of conventional tryptic DIA discovery proteomics through split digest, pooled C18 cleanup, single-shot DIA acquisition, and unified downstream analysis in the mE-DIA Pipeline, powered by DIA-NN (Demichev et al., 2018). We benchmarked the computational workflow against DDA-dependent analyses in human cell-line proteomes and demonstrate the integrated workflow’s applicability to profile start/stop sequence proteoforms (i.e., protein species with non-canonical N-termini (start) or premature, non-canonical C-termini (stop) relative to the annotated sequence) in two bacterial pathogens. The toolset includes the mE-DIA Pipeline graphical user interface and the mE-DIA Peptide Mapper viewer for browsing results.

Together these components extend discovery DIA proteomics by the ability to detect and quantify start/stop proteoforms inaccessible to conventional single-enzyme and gene-centric analysis.

## Results

### The Library-free mE-DIA Workflow

The mE-DIA workflow integrates separate multi-enzyme proteome digestion with combined DIA-MS acquisition, library-free analysis and proteoform inference into a single, accessible analytical framework (Fig. 1). In the experimental stage, protein samples are digested in parallel using complementary proteases and combined into one peptide mixture prior to peptide purification or prior to a single DIA-MS injection from which rich peptide level evidence is extracted by enzyme de-multiplexing analysis in the mE-DIA Pipeline (Fig. 1A). This experimental design represents a minimal deviation from the conventional tryptic digestion workflow and enables the simultaneous recording of MS/MS ions across extended sequence spaces. On the computational side (Fig. 1B), the workflow entails two modules. In the first module, a 2-pass multi-enzyme search combines per-enzyme reference protein database digestion, *in silico* spectral library prediction and pass 1 search of the DIA dataset to produce per-enzyme empirically corrected spectral libraries. These libraries are then merged for concurrent query in combined pass 2 analysis, producing the enriched multi-enzyme quantitative data on precursor, peptide and protein group level under one cumulative q-value and error estimation (Fig. 1B, upper panel). Optionally, the analysis can be extended to query semi-specific peptide species conforming to the detected protein background (Fig. 1B, lower panel). Here, the confidently identified protein groups recovered during the fully specific pass 2 analysis are used to constrain the otherwise prohibitively large semi-specific search space. For these proteins, enzyme-specific semi-specific *in silico* digestion and spectral library prediction are performed, followed by pass 3 query of the DIA dataset generating empirically refined semi-specific spectral libraries. In the final analysis stage, these semi-specific libraries are combined with the previously established pass 1 empirical libraries for integrated pass 4 analysis, producing unified quantitative matrices across precursor, peptide, and protein-group levels, substantially extending peptide-level evidence with non-canonical, semi-specific peptide sequences. Finally, a proteoform inference step identifies candidate start/stop sequence proteoforms, i.e. non-canonical N-or C-terminal backbone boundaries relative to the annotated sequence. Each backbone start or stop position at a given residue that is evidenced by two or more distinct peptide sequences is reported as a *proteoform-differentiating event*, with a proteoform identifier and a quantitative matrix of the supporting peptide precursor signals across the study cohort reported for further interrogation. We provide an mE-DIA Pipeline graphical user interface that drives setup, execution and visualization of multi-enzyme DIA analyses (Fig. S1A) and an additional mE-DIA Peptide Mapper tool to utilize proteoform level insights (Fig. 1C). Specifically, the mE-DIA Peptide Mapper provides interactive visualization of peptide evidence along all detected protein’s backbones in a condition-or run-dependent manner, as well as a differential discovery module to prioritize proteins for review that display conditional differences in amino acid observation frequency along their backbone, compared across user-defined groups (for details, see Methods section). Together, the framework provides an end-to-end solution from raw DIA data to peptide mapping and discovery of proteoform-level insights, enabling researchers to effectively extract and navigate proteoform-level discordance evident along the different states of the experimental system of interest.

**Figure 1:**
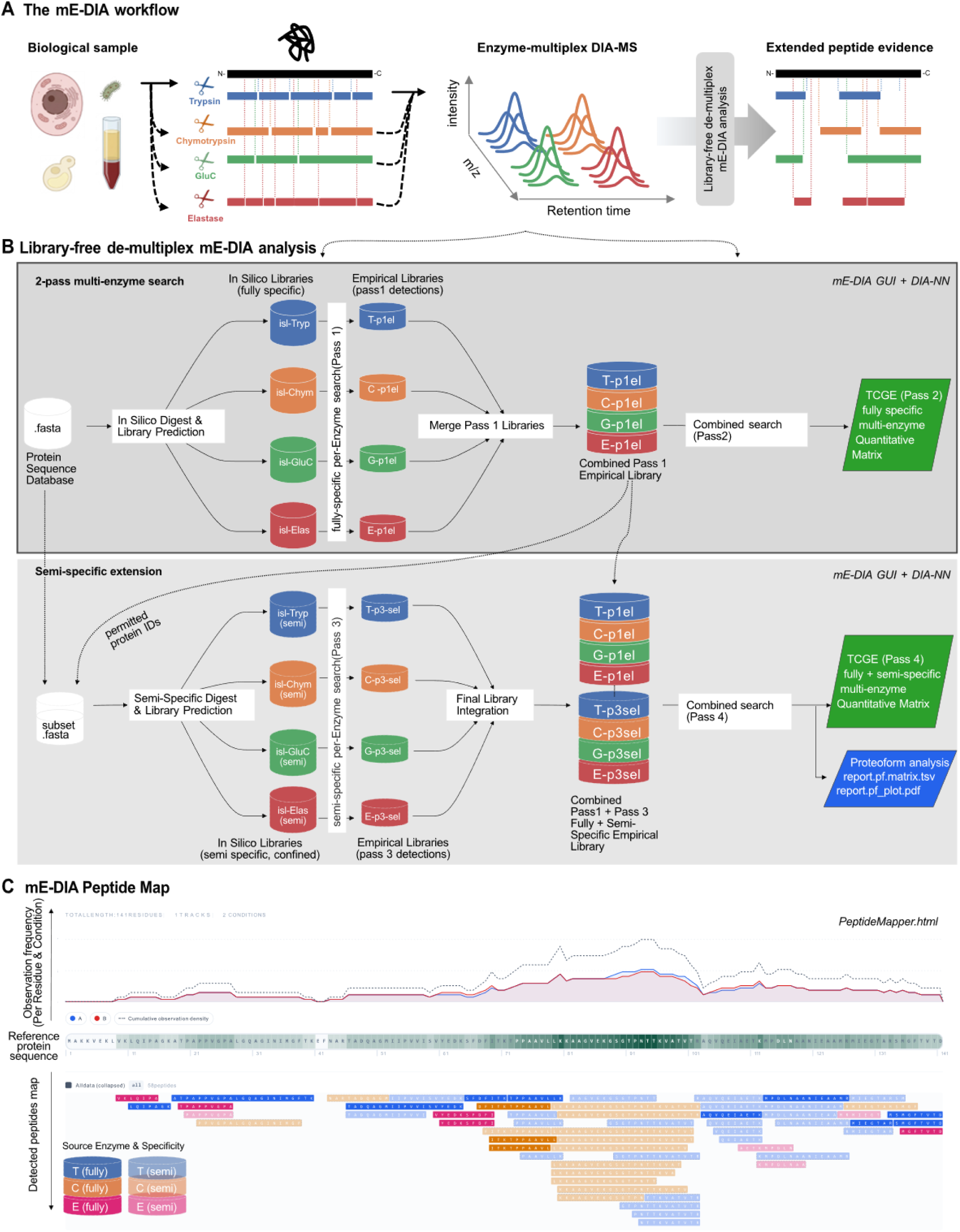
The mE-DIA experimental s computational workflow. **(A)** The overall mE-DIA workflow scheme, including experimental section. Protein samples from diverse biological matrices are digested separately with complementary proteases and generated peptide sets pooled into a single peptide mixture and acquired in one enzyme-multiplex DIA-MS injection, recording distinct peptide populations and co-eluting fragment ion patterns in a shared retention-time and m/z grid and DIA data file. The peptides across the different sequence spaces are subsequently extracted and quantified in per-enzyme peptide-centric analysis and integration in library-free de-multiplex mE-DIA analysis, yielding extended and orthogonal peptide evidence along detectable protein backbones, with semi-specific peptide sets delineating candidate protein backbone variants through the observation of non-canonical N- or C-termini. **(B)** Computational flow scheme of library-free de-multiplex mE-DIA analysis. Top: 2-pass multi-enzyme search. The reference protein sequence database is digested and peptide spectral libraries predicted in silico per enzyme and in silico (isl-) libraries queried against the DIA data in per-enzyme pass-1 searches to derive empirically corrected libraries. Per-enzyme pass-1 libraries are subsequently merged into one combined pass-1 empirical library which is then queried against the same DIA data in pass 2 to produce a unified fully specific multi-enzyme quantitative matrix. Bottom: Semi-specific extension. Protein groups confidently identified in pass 2 define a subset FASTA, from which per-enzyme semi-specific in silico libraries are predicted and queried in pass 3. The resulting empirical semi-specific pass-3 libraries are combined into one combined library spanning all enzyme specificities and including also the pass-1 fully specific library for the final, integrated pass-4 search, yielding the fully and semi-specific multi-enzyme quantitative matrix and triggering the extended proteoform inference module and -outputs. **(C)** Example mE-DIA peptide map from the interactive, multi-track mE-DIA Peptide Mapper tool built to visualize and interrogate multi-enzyme and candidate protein variant results. The main peptide map interactively illustrates multi-enzyme fully- and semi-specific peptide observations along reference protein backbones (bottom) as well as conditional residue observation frequency profiling (top) that underlies the Differential Discovery module (not shown). The mE-DIA Pipeline graphical user interface to configure and monitor mE-DIA analyses is shown in Fig. S1A. See main text and Methods for details.

### Comparing Library-free mE-DIA data analysis to state-of-the-art DDA-based-workflow

To contextualize the mE-DIA library-free data analysis module against the data processing workflow of the previously established implementation of multi-enzyme DIA proteomics, we reanalyzed a subset of the raw enzyme-multiplex DIA data generated by Richards *et al*. (from dataset PXD027242), in which lysates of the human cell lines HSC-6 and SCC-25 were digested separately with trypsin, Glu-C and Asp-N prior to pooling and label-free acquisition in a single DIA injection per replicate ( (Richards et al., 2022), design of the comparison in Fig. 2A). In the original implementation, the individual single-enzyme digests were additionally measured in DDA-MS mode to derive enzyme-specific spectral libraries that were subsequently merged into one library used to query the different peptide sets in the enzyme-multiplexed DIA runs, whereas the library-free re-analysis via mE-DIA relied solely on the DIA injections (Richards et al., 2022).

**Figure 2:**
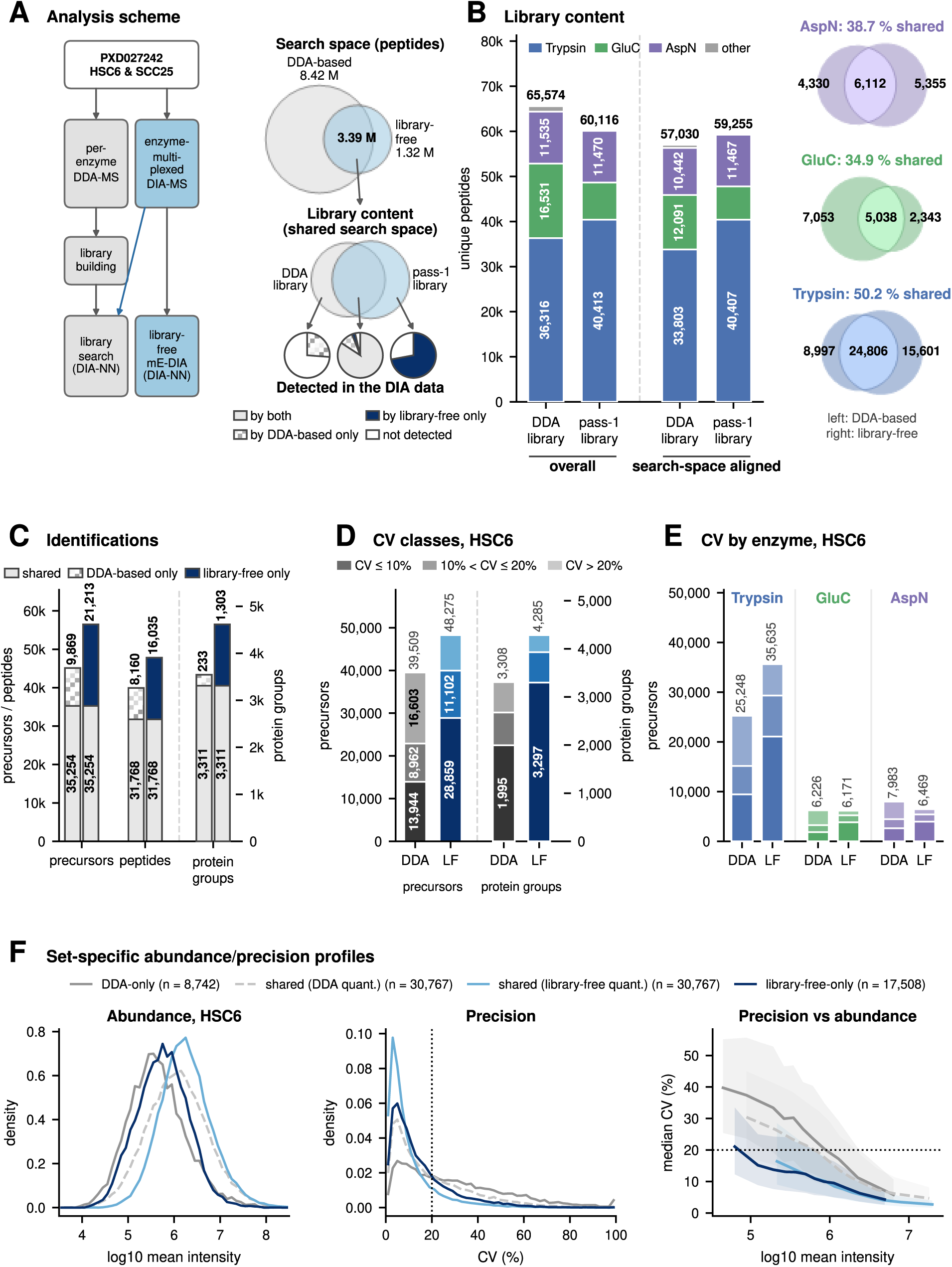
Benchmarking library-free mE-DIA analysis against the DDA-library-based workflow on PXD027242 human cell line multi-enzyme DIA data. (A) Study design (left) and analyte flow (right) illustrating the scope of the comparison between the two analysis workflows. Within their shared search space, library content and final set of analytes detected and quantified consistently in the DIA data. (B) Library content: unique modified peptides per enzyme in the DDA library and the pass-1 library, overall and search-space aligned (restricted to the shared search space). Venn diagrams (right): Per-enzyme overlap of distinct peptides between the two search-space aligned libraries, DDA-based left and library-free right. **(C–F)** Analytes within the shared search space. **(C)** Identifications at precursor, peptide (modified sequence) and protein-group level, shared and workflow-exclusive, counting analytes identified in all three replicates of a cell line at 1 % experiment-wide FDR, unioned across the two cell lines. **(D)** Quantitative precision in HSC6: precursors (left axis) and protein groups (right axis) binned by CV across the three replicates into ≤ 10 %, 10–20 % and > 20 %; numbers above bars, analytes in each workflow’s matrix; protein-group quantities recomputed from the retained precursors for both workflows. **(E)** Precursor CV classes by enzyme, HSC6. **(F)** Set-resolved analysis of precursor abundance (left), precision distribution (middle) and binned median CV with interquartile band versus abundance (right) of precursors reported by one or both workflows, HSC6; shared precursors are shown with each workflow’s own quantification.

To control for differences between pipeline’s search space due to divergent parameterization, and owed to hardware limits preventing to fully mirror the DDA pipeline’s settings on the DIA side, we based the comparison exclusively on the peptide analytes from the shared search space of both pipelines, i.e. peptides that both workflows searched for in the MS data, and follow their representation in the library and final detectability in the DIA data (Fig. 2A, for details, see Methods). The DDA-based search space was significantly larger, probing ∼ 12 M peptidoforms including variably oxidized methionine or acetylated protein N-termini, peptides longer than 30 aa and > 2 missed cleavages, compared to the library-free pipelines’ overall narrower, 4.7 M peptidoform-spanning search space. The following comparisons are limited to observations of the ∼3.4 M peptidoforms that both pipelines queried in the data.

First, we compared the contents of the enzyme-specific-spectral library Richards et al. generated from the duplicate DDA-MS measurements per each enzyme digest through a multi-step pipeline involving MaxQuant, SpectraST, and Spectronaut to the contents of the library derived directly from their enzyme-multiplex DIA data recorded in triplicate across the HSC6 and SCC25 cell line’s multi-enzyme digest pools, as generated by mE-DIA pass 1 library-free analysis (‘Combined Pass 1 Empirical Library’, compare Fig. 1B, upper panel & Fig. 2 ACB). Overall, the deposited DDA library contained 65,574 unique peptides, ∼ 9 % more than the library-free pass-1 library of 60,116 unique peptides, driven by markedly deeper coverage of Glu-C-derived peptides overcompensating for less tryptic peptides (Figure 2B, left). When restricted to peptides originating from the shared search space, the DIA library shows deeper coverage in terms of total absolute numbers with 59,255 vs 57,030 unique peptides, with an enzyme-specific yield of 40,407 vs 33,803 tryptic, 7,381 vs 12,091 Glu-C-derived and 11,467 vs 10,442 Asp-N-derived peptides in DIA pass1 vs. the DDA library, respectively. Despite the shared search space control, the two libraries converge strikingly little: 50 %, 35 % and 39 % of the distinct peptide sequences in either search-space aligned library are present in both, for trypsin, Glu-C and Asp-N respectively (Fig. 2B, right panel). Different observability profiles are expectable in split per-enzyme DDA vs. enzyme-multiplex DIA acquisition and different algorithmic paths to the library, and the impact of either not accessible from this comparison alone.

More importantly, we next evaluated which of those peptides were ultimately detectable and quantifiable in the DIA data. Considering only analytes within the shared search space and that were identified consistently in three out of three replicates in at least one of the two cell lines (Fig. 2C), the library-free mE-DIA analysis identified more analytes than the DDA-library analysis at every level: 56,467 versus 45,123 precursors (+25.1 %), 47,803 versus 39,928 peptides (+19.7 %) and 4,614 versus 3,544 protein groups (+30.2 %) (Fig. 2C). The workflows provide access to partly orthogonal analyte sets: DDA-based analysis exclusively retrieved 8,160 peptides and 233 protein groups that were searched for but out of reach under the library-free paradigm, whereas library-free mE-DIA exclusively detected 16,035 peptides and 1,303 protein groups that were inaccessible via the DDA-library route (Fig. 2C). As expected, recall of the 17,209 peptides only in the DDA library was lower (27 %) than recall of the 20,870 peptides only in the pass-1 library (70 %) (Fig. 2A), in line with DDA sampling peptides in single-enzyme runs that are undetectable in the enzyme-multiplexed DIA runs, and with library-free preferential sampling of detectable peptides under the matched acquisition conditions by design. It is notable that 4,615 peptides were accessed only through the DDA route, providing evidence of per-enzyme sampling retrieving peptide evidence that is consistently observable in the DIA data but that is missed in the library-free route. Each workflow thus provides access to analytes in the DIA data that the other does not report, with an overall lead for the library-free route. On the level of detected protein groups 3,311 were common to both workflows, covering 93.4 % of the DDA-based result. For the shared protein groups, library-free analysis yielded a median of eight peptide sequences, compared to six in the DDA-based result. Workflow-exclusive protein groups rested on less peptide level evidence on either side, with a median of two and one peptide sequences in the library-free and DDA-based workflow, respectively. Of the uniquely identified groups, 561 of 1,303 (43.1 %) and 176 of 233 (75.5 %) were identified based on a single peptide sequence. For the library-free workflow, we cannot exclude that part of this margin reflects identification leakage under non-standard application of tools developed for single-enzyme use and therefore examined the error behavior of the multi-enzyme library-free workflow directly in a dedicated leave-one-out experiment (See below and Fig. 5).

While coverage is a key metric, quantitative reliability is a proxy for a workflow’s potential for biological discovery and thus provides an additional dimension to further classify workflow-dependent observations, leading us to interrogate analytical precision of both workflows among the observed peptide precursors and protein groups within the shared search space (Fig. 2D–F). The library-free workflow returns more analytes quantified at high precision. Considering analytes quantified at ≤ 20 % CV, the library-free workflow yields 39,961 versus 22,906 precursors compared to the DDA-based workflow, a gain of +74 % (HSC6 cell line samples, Fig. 2D). At protein-group level, library-free analysis yields 47 % more protein groups quantified at ≤ 20 % CV (3,930 versus 2,677 in HSC6; 3,871 versus 2,623, +48 %, in SCC25), compared with a +29 % gain at identification level (4,285 versus 3,318 in HSC6). Stratifying precursors by enzyme (Fig. 2E) reveals that the library-free workflow’s gain in precursor identifications was driven exclusively by tryptic peptides, with fewer Glu-C-derived (6,171 versus 6,226) and Asp-N-derived (6,469 versus 7,983) precursors (Fig. 2E). However, all three enzyme channels see a marked increase of well-quantifiable precursors (≤ 20 % CV, +93, +61 and +21 % for Trypsin, Glu-C and Asp-N, respectively) (Fig. 2E). Both workflow-specific precursor sets were of overall lower abundance than the shared set, yet span well into the higher abundance range (Fig. 2F, left panel). Library-free exclusive precursors were quantified more precisely than those recovered exclusively by the DDA-based workflow (median CV 10.6 % vs. 25.0 %, Fig. 2F, middle panel). Notably, also the shared precursors were quantified more precisely by the library-free workflow (median CV 6.6 % vs. 13.8 %, Fig. 2F, right panel), consistently throughout the abundance range, with all precursor sets following the typical trend of increased variability at lower abundances (Fig. 2F, right panel). These results show that the integrated library-free workflow on the same data provides overall higher quantitative reliability and scope and statistical power for discovering biological variation and that the per-enzyme DDA and spectral library-based workflow might suffer from divergent sample and acquisition conditions that lead to the selection of query parameters that are suboptimal to quantify peptides from enzyme-multiplexed DIA maps. Without acquiring dedicated single-enzyme DDA runs or building an experimental spectral library, library-free mE-DIA delivers deeper identification and higher quantitative fidelity on the same raw data.

### Extended evidence basis for bacterial pathogen proteoform landscape

To explore the analytical performance of the multi-enzyme DIA workflow under controlled conditions, we analyzed the proteomes and proteoform landscapes of the two bacterial pathogens *Streptococcus pyogenes* strain SF370 and *Escherichia coli* strain O18K1.

First, we analyzed *S. pyogenes* using a three-enzyme TCE multiplex (trypsin, <u>c</u>hymotrypsin, <u>e</u>lastase) alongside matched single-enzyme DIA runs to quantify the impact of multiplex analysis. In fully specific (pass 2) mode, the single-enzyme runs yielded 19,820, 12,679 and 8,726 precursors for trypsin, chymotrypsin and elastase, respectively (14,366 / 9,743 / 7,303 peptides; 1,294 / 1,170 / 925 protein groups), with trypsin producing the most identifications, followed by chymotrypsin and then elastase (Fig. 3A, solid bars). The TCE multiplex produced 21,246 precursors (17,190 peptides; 1,198 protein groups) within a single injection, i.e. 1.07-fold the precursors and 1.20-fold the peptides of the best single-enzyme run, but 0.93-fold its protein groups (Fig. 3A–B). Individual and multiplex acquisition largely converged on the same set of proteins in the sample, albeit with individual-enzyme-specific proteins inaccessible through the multiplex workflow (85, 32 and 4 protein groups exclusive to individual trypsin, chymotrypsin and elastase analysis, respectively, Fig. 3C), whereas 11 protein groups only passed confidence thresholds in the combined, multiplex workflow (Fig. 3C).

**Figure 3:**
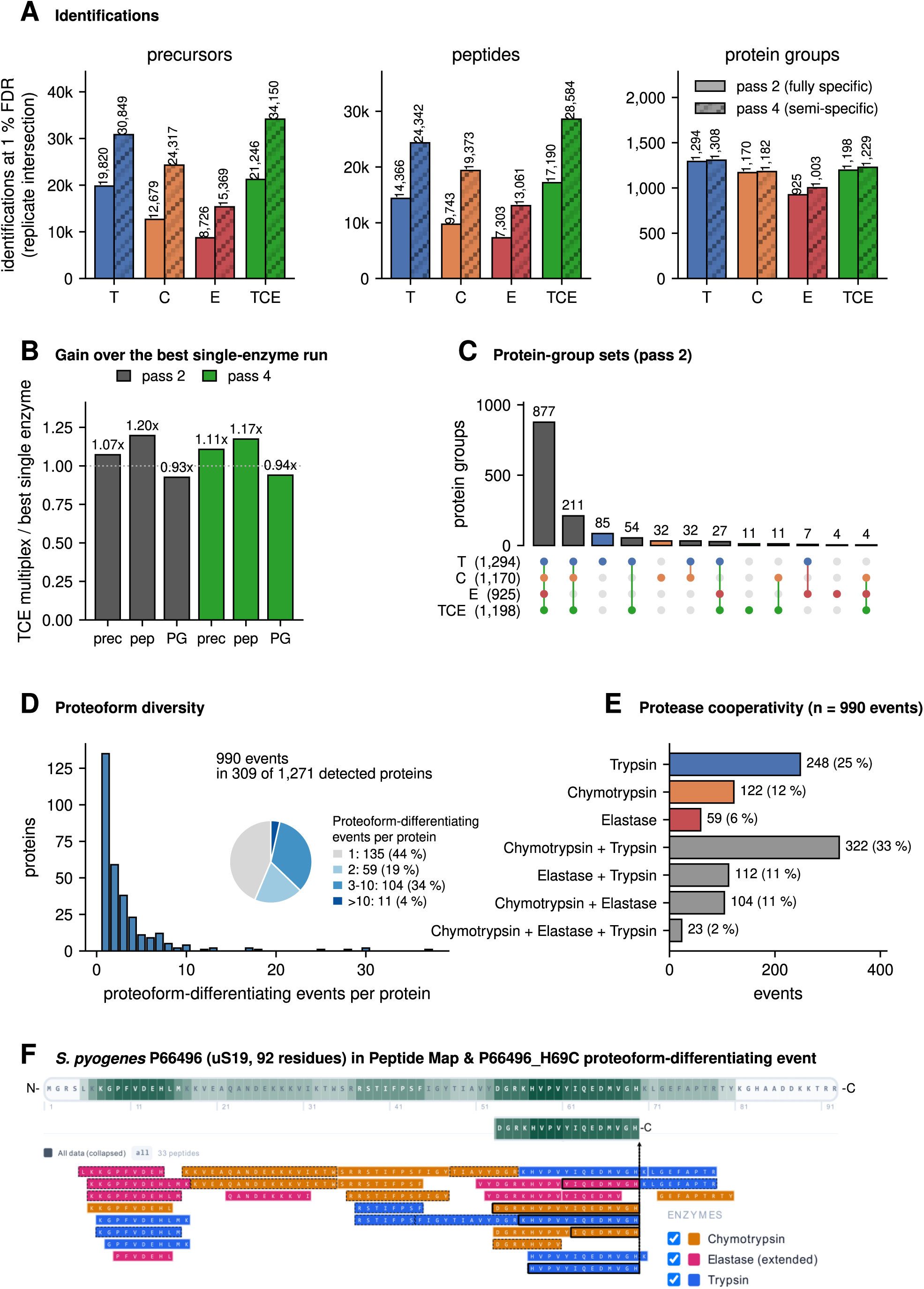
Multi-enzyme DIA analysis of *S. pyogenes* SF370 with the TCE (Trypsin / Chymotrypsin / Elastase) multiplex. **(A)** Precursor, peptide and protein-group identification counts across individual single-enzyme and TCE multiplex analyses in fully specific (pass 2) and semi-specific extended (pass 4) modes. **(B)** Relative gains of the TCE multiplex over the best single-enzyme run, in both pass 2 and pass 4. **(C)** UpSet plot of protein-group overlap between individual T/C/E and TCE multiplex (pass 2). **(D)** Distribution of proteoform diversity per reference protein in the TCE multiplex (990 events in 309 of 1,271 detected proteins, after removal of ISD products). **(E)** Per-enzyme and per-enzyme-combination contribution of peptide evidence to those events. **(F)** Peptide Mapper view of P66496 (30S ribosomal protein uS19) in the TCE multiplex with fully and semi-specific peptides, where the H69 C-terminal proteoform is resolved (H69C terminus indicated by the black vertical arrow); the supporting peptide sequences are marked by black boxes and supporting precursors are listed in Fig. S1B. Per-enzyme recall within the multiplex and channel-restricted protein-group retrieval are given in Fig. S3, and the proteoform inference summary in Fig. S5.

The semi-specific extension (passes 3 and 4) forms the basis for proteoform discovery by adding substantial sequence space in all conditions (Fig. 3A & Fig. S4A). Semi-specific analysis added +92 % precursors for individual chymotrypsin, +76 % for elastase and +56 % for trypsin, consistent with a higher prevalence of semi-specific cleavage products for the non-tryptic enzymes and a substantial number of analytes not accessed via the conventional, fully specific search parameters. Within the TCE multiplex, the semi-specific pass yielded +61 % additional precursors and +66 % additional peptides, reaching 34,150 precursors and 28,584 peptides in total, with gains per enzyme level mirroring the gains seen in single-enzyme analysis (Fig. 3A & Fig. S4A). Anchored on proteins identified in pass 2, the semi-specific extension incurred only minor changes in protein groups the detectable peptide set resolved to, with changes ranging between +2.6 % in the TCE multiplex and +1.0 % to +8.4 % for the individual enzymes, i.e. the semi-specific gain is, by design, additional sequence-level evidence for an essentially unchanged set of proteins rather than additional proteome depth.

Peptide length, charge state, chromatographic peak width and retention-time prediction error per enzyme, for all three datasets, are shown in Fig. S2. Elastase released the shortest peptides (mean 9.2–9.4 residues) at the lowest charge states (mean 1.95–1.98), chymotrypsin was intermediate (11.7–11.8 residues), and Glu-C and Asp-N peptides resembled tryptic ones (12.8–13.6 against 12.8–13.2 residues), while peak widths were indistinguishable between enzyme channels within a dataset (Fig. S2).

Having established the analytical performance of the multi-enzyme workflow, we next explored what the expanded peptide evidence reveals about protein sequence diversity. Peptides carrying N- or C-termini not attributable to the digestion enzymes can provide evidence of non-canonical protein termini and candidate proteoforms. Using at least two distinct peptide sequences to support each terminus, and after removing ISD (in-source decay) products, i.e. truncated peptides formed in the ion source from a co-eluting, more intense longer peptide (Kim et al., 2013; Schramm et al., 2026), the TCE multiplex identified 990 proteoform-differentiating events across 309 proteins (Fig. S5A, B). The ISD filter flagged 142 supporting precursors and disqualified 75 of 1,065 candidate events, 74 of them N-terminal (Methods, Fig. S5D). The bonds cleaved in the flagged products most often follow V, A or I (20 %, 15 % and 11 % of 362 fragment–parent pairs across both organisms), the residue preference reported for in-source fragmentation (Schramm et al., 2026). This represents a 5.3-fold increase over the 186 events identified in the conventional trypsin-based workflow (122 proteins), whose N-terminal candidates were most affected by the ISD filter (75 of 176 removed), and 1.8-fold the 551 events identified by the most productive single-enzyme digest, elastase (268 proteins). Overall, the TCE multi-enzyme strategy identified 2–38 differentiable proteoforms per protein, arising from 1–37 sequence divergence events per protein (termination in the C-terminus or alternate start in the N-terminus, each counted as one proteoform), with each event evidenced by 2–5 independently detected peptide sequences (Fig. 3D). Because the peptides sharing a terminus were released by different proteases, and the digests were quenched before they were combined (Methods), that terminus cannot have been generated by any one of them: 561 of the 990 events (57 %) are evidenced by peptides from at least two of the three proteases and 23 by all three (Fig. S5C). Restricting the evidence to one enzyme channel at a time, under the same two-sequence rule, calls 277, 151 and 86 events from trypsin, chymotrypsin and elastase peptides respectively; 514 in total, against 990 from the combined evidence (Fig. S5).

Roughly half the events therefore exist only because peptides released by different proteases coincide on the same terminus, and are not reachable from any single enzyme channel of the multiplex. The analysis is performed on the full study and associates the peptide precursors with the proteoform they evidence, extracting the respective quantitative data for discovery of changes between experimental conditions. To illustrate how different enzymes cooperate in evidencing non-canonical sequence variants, we decomposed the peptide evidence base of the 990 sequence divergence events by contribution of peptides generated by respective enzyme, identifying trypsin+chymotrypsin as the most productive enzyme pair, jointly evidencing 322 of the 990 events (33 %), and suggesting that pair as an efficient two-enzyme split digest system to balance peptide load, interference and proteoform discovery (Fig. 3E).

As an example proteoform-differentiating event, we identified the P66496_H69C variant with a C-terminus at histidine 69 of the *S. pyogenes* small ribosomal subunit protein uS19 (30S ribosomal protein S19). This variant was not called in the individual trypsin analysis, yet in the multiplex is evidenced by five peptide sequences and eight precursors, none of them flagged as ISD products (Fig. 3F, Fig. S1B). The five sequences were released by three different proteases, trypsin, chymotrypsin and elastase, and all share the H69 C-terminus while their opposite termini remain enzyme-specific, so the truncation pre-dates the digest and is most consistent with endogenous processing rather than a digestion artefact. Of the individual analyses, the chymotrypsin analysis also calls the event, from chymotryptic peptides alone, albeit without the added certainty of cross-enzyme observability. Two further peptides span H69 rather than ending at it, indicative of the presence of a long protein variant in parallel to the short variant, in the same injection and sample. The example demonstrates how orthogonal peptide classes generated by complementary proteases can synergistically expose non-canonical proteoforms that would otherwise evade detection due to enzyme-specific peptide blind spots, and how this is enabled by the mE-DIA Pipeline, its proteoform analysis and the Peptide Mapper tool. Dedicated follow-up studies may now explore the biological function and therapeutic target potential of such newly resolved proteoform structures.

To isolate the effect of pooling digests on a technical level, we compared each single-enzyme analysis with the same enzyme’s channel of the multiplex with 1/3^rd^ peptide load per enzyme (fully specific pass 2 on both sides; Fig. S3). The multiplex recalled 66 % of the peptides identified in the trypsin analysis, 49 % for chymotrypsin and 33 % for elastase, Fig. S3A). At protein-group level the multiplex retains almost the full population, recalling 90 %, 94 % and 98 % of the groups of the trypsin, chymotrypsin and elastase analyses (Fig. S3C). Restricted to the peptides of the respective enzyme channel, recall falls to 89 %, 78 % and 70 % (Fig. S3C), so the other channels supplied the evidence for groups a weak channel lost. Quantitative precision was not compromised in the multiplex, with precursor-level replicate correlation stable between multiplex and the individual analyses (Fig. S4B, C). To weigh what pooling returns against what it costs, we combined the three single-enzyme analysis results into a combined, “pseudo-pool” result set and passed it through the same proteoform inference (Fig. S4D). With three times the acquisition time and sample, the separate acquisition with pseudo-pooling yielded 2,386 candidate proteoform events against 990 for the multiplex, as well as twice the peptides (56,097 vs 28,584). Per hour of acquisition, however, the multiplex delivers 1.2-fold more candidate events (248 vs 199) and 1.5-fold more peptides (7,146 vs 4,675), at 90 % of the combined pseudo-pool’s protein groups (1,229 of 1,364).

These results do show attrition through multiplexing, likely owed to three-fold lower peptide load moving lower-abundance peptides below the limit of detection in line with recall declining with the depth of the individual channel; and losses possibly due to interference by and charge competition with co-eluting peptides generated by the other enzymes. However, losses are non-linear and enzyme multiplexing appears as an efficient choice when instrument time or sample, rather than depth for an individual protein or enzyme, limits the analysis.

In addition to the three-enzyme analysis of *S. pyogenes,* we applied the workflow to an *Escherichia coli* isolate (strain O18:K1), using four-enzymes (trypsin, chymotrypsin, elastase and Glu-C). While these runs were performed primarily to support the evaluation of error rates in the leave-one-out-analysis (See below & Fig. 5), the 4-enzyme-multiplex also allowed us to survey proteoform differentiating events observable in this second gram-negative pathogen (Fig. 4A-C). Overall, 1,564 reference protein groups were identified based on 28,083 distinct peptide sequences and 33,531 precursor ions in the semi-specific pass 4 (Fig. 4A). For 250 of the reference proteins, 544 proteoform sequence variants were identified, with diversity spanning 2–18 differentiable proteoforms per reference protein, each evidenced by at least two independently detected peptide sequences, each in turn evidenced by one or more peptide precursor ion signal (Fig. 4C). As in *S. pyogenes*, most events rest on more than one protease: 335 of the 544 (62 %) are evidenced by peptides from at least two of the four enzymes, 28 by three (Fig. S5C). Again, trypsin and chymotrypsin combined showed to synergize most effectively in cooperative proteoform identification, with the newly added Glu-C contributing peptide evidence at 92 of the 544 sites (17 %), the smallest share of the four enzymes (chymotrypsin 329 sites, 60 %; trypsin 291, 53 %; elastase 195, 36 %) (Fig. 4D).

**Figure 4:**
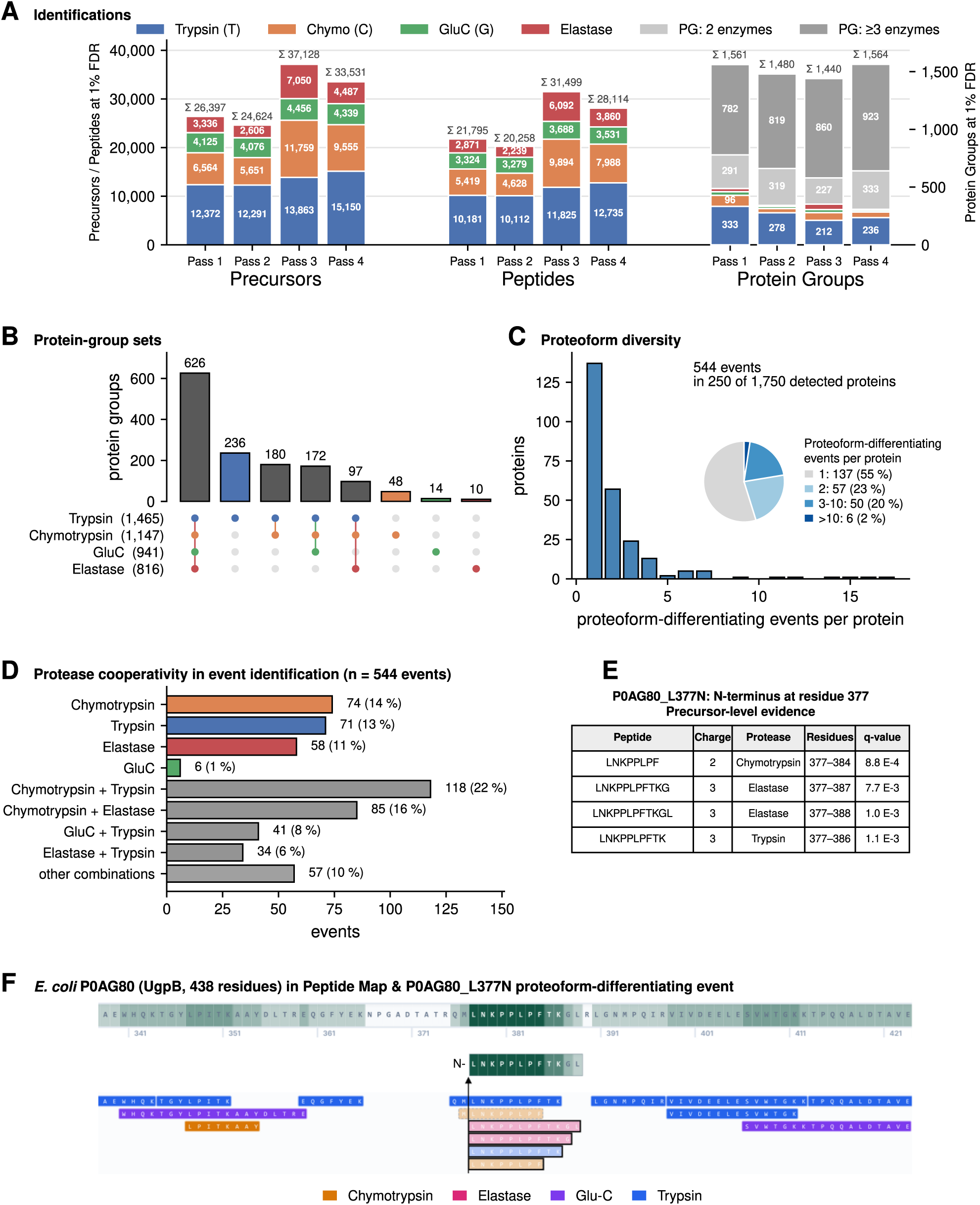
Four-enzyme mE-DIA analysis of an *Escherichia coli* isolate (O18:K1). **(A)** Precursor, peptide and protein-group identification counts across the four mE-DIA passes (Pass 1 empirical, Pass 2 combined fully specific, Pass 3 semi-specific, Pass 4 semi + combined), stacked by enzyme channel (Trypsin, Chymotrypsin, Glu-C, Elastase) and, for protein groups, by the number of enzymes contributing supporting evidence. **(B)** UpSet plot of protein-group set membership across the four enzyme channels in pass 4, showing the five largest sets and every set unique to one enzyme. **(C)** Distribution of proteoform diversity per reference protein (544 sequence-divergence events in 250 of 1,750 detected reference proteins, after removal of ISD products). **(D)** Per-enzyme and per-enzyme-combination contribution of peptide evidence to proteoform sequence-divergence event identification. **(E)** The peptide precursors supporting the P0AG80_L377N call, with charge, protease, residue range and experiment-wide precursor q-value, and beneath the table the peptides that span residue 377 rather than starting at it, which evidence the co-existing full-length form. **(F)** Peptide Mapper view of P0AG80 (UgpB, 438 residues; excerpt of residues 340–422 shown) in the TCGE multiplex, with peptides coloured by protease. The truncated proteoform is drawn from L377 to residue 388, the extent covered by its supporting peptides, whose sequences are outlined in black; the new N-terminus is indicated by a vertical arrow, and the precursor evidence is listed in panel **(E).**

A comparable example proteoform-differentiating event in this dataset is P0AG80_L377N in the periplasmic sn-glycerol-3-phosphate-binding protein UgpB (438 residues), where an N-terminus at leucine 377 is evidenced by four peptide sequences and four precursors released by three different proteases: chymotrypsin (residues 377–384), trypsin (377–386) and elastase (377–387 and 377–388), whose C-termini are enzyme-specific while their N-termini coincide at L377 (Fig. 4E, F). All four precursors pass 1 % experiment-wide FDR (q = 8.8 × 10⁻⁴ to 7.7 × 10⁻³), and none is flagged as a suspect ISD product. The preceding bond, M376–L377, is not a cleavage site under the search rule of any of the four enzymes. Chymotrypsin can cleave after methionine as a minor activity, but the tryptic and elastase peptides that share this N-terminus were released in separate digests that were quenched before pooling, so the terminus pre-dates the digest. Two further peptides span residue 377, a fully specific tryptic peptide (375–386) and a semi-specific chymotryptic peptide (376–384), in line with a long variant, potentially full-length, protein being present in the same sample and the truncated species co-exists with it.

### Robustness of multi-enzyme DIA assessed through Leave-One-Out Validation

To evaluate the robustness of the enzyme-multiplexed analysis workflow and assess the risk of cross-enzyme false identifications, we implemented a leave-one-out validation (LooV) design in which each enzyme was omitted from one of the samples. As a proxy for error rates, we assessed leakage rate of analytes not contained in the sample but spuriously matched and reported by the analysis framework at two levels.

First, the leakage rate in combined pass 2 analysis was assessed by retaining all false target peptides in the composite pass 1 empirical library, including those that are no longer contained in the left-one-out sample. A precursor counts as a false identification when the library attributes it exclusively to the omitted enzyme; peptides that a present enzyme can also generate are not counted. In this worst-case scenario where all targets of the given peptide class are false, default q-value thresholding at ≤ 0.01 yielded precursor-level leakage of 1.7, 1.3, 0.5 and 0.6 % of total results and 1.8, 3.9, 2.1 and 4.1 % leakage as fraction of library-contained false targets for trypsin, chymotrypsin, Glu-C and elastase peptide sets, respectively. On protein level (thresholded on Global.PG.Q.Value), where a protein group counts as false only when all its precursors are false, raw leakage reached 0.6, 0.9, 0.6 and 0.1 % of results and 1.5, 7.5, 13.6 and 1.7 % leakage rate from false targets in library, respectively (Fig. 5A). Leakage as a function of the confidence threshold, rather than at the default cutoff alone, is shown in Fig. 5B.

**Figure 5:**
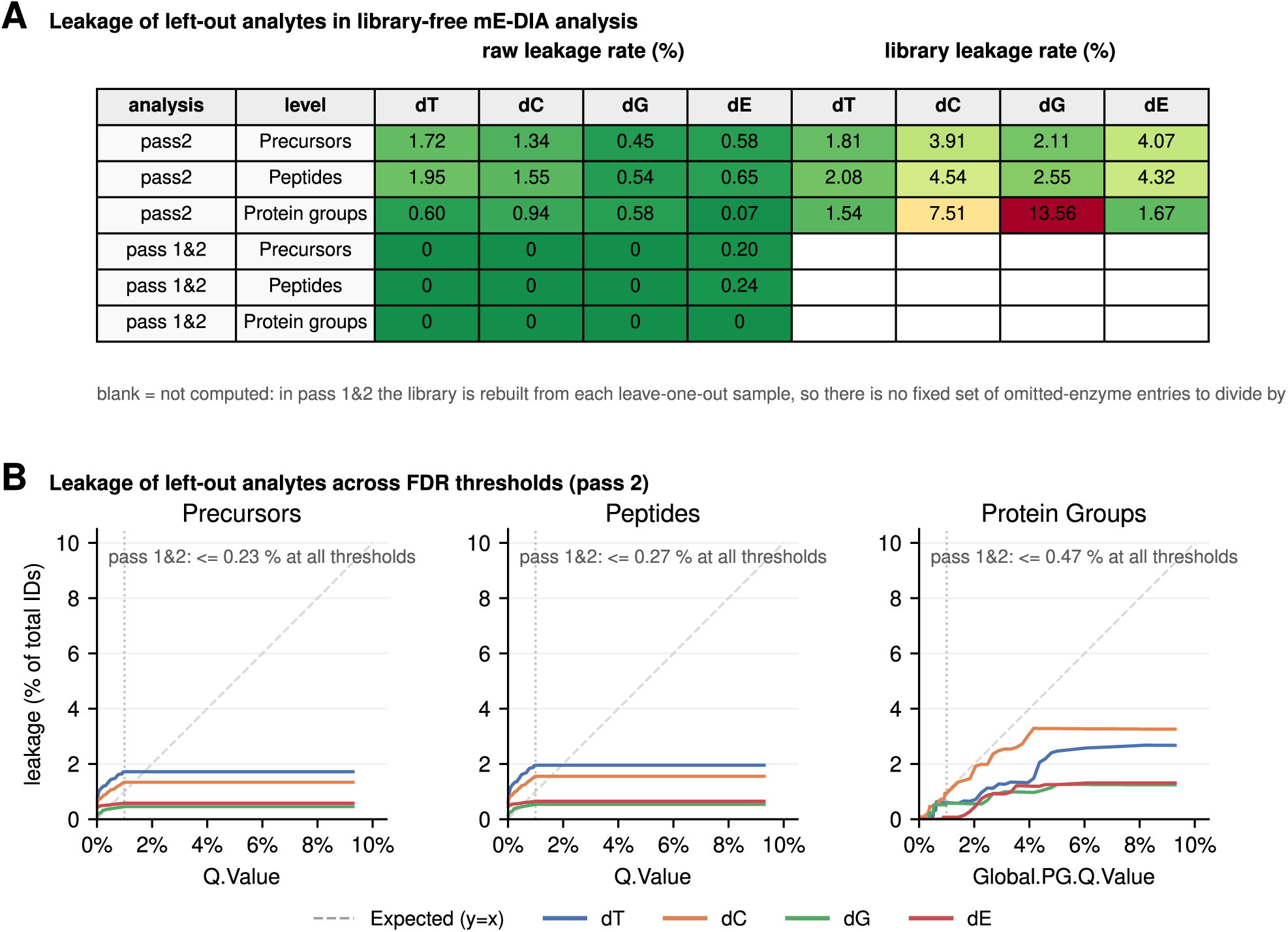
Leave-one-out validation (LooV) of cross-enzyme error control in the four-enzyme *E. coli* multiplex. **(A)** Error estimation through analyte leakage in LooV analysis summary table. For each omitted enzyme (dT, dC, dG, dE), raw leakage (fraction of total reported IDs originating from the omitted enzyme channel) and library leakage (fraction of omitted-enzyme entries in the library reported as IDs) at precursor, peptide and protein-group level, in pass-2 mode (composite library retained) and pass-1C2 mode (library rebuilt from the LooV sample). Library leakage is not computed in pass-1C2 mode, where the library is rebuilt from each leave-one-out sample; those cells are left blank rather than reported as zero. Wrong identifications are precursors attributed exclusively to the omitted enzyme. **(B)** Leakage as a function of the Q.Value (precursor, peptide) or Global.PG.Q.Value (protein-group) threshold for each omitted enzyme in pass-2 mode. Default 1 % FDR cutoff indicated by the vertical line. Note that in the complete pass-1C2 workflow curves stayed at 0 or below 0.5 % across the threshold range and their curves are therefore not drawn. Instead, maxima are stated as inserts in the panel. See main text and Methods for details.

Secondly, the realistic two-pass scenario was evaluated by rebuilding the empirical library from each LooV sample (pass 1) before performing the combined pass 2 search. Under these conditions, raw leakage fell to zero for the trypsin, chymotrypsin and Glu-C omissions at every analyte level, and to 48 of 23,776 precursors (0.20 %) and 46 of 19,442 peptides (0.24 %), with no protein group, for the elastase omission (Fig. 5A). The library leakage rate is not computed in this mode, because the library is rebuilt from each leave-one-out sample and therefore holds no fixed set of omitted-enzyme entries to serve as its denominator. That the rebuilt libraries leave little to leak plausibly reflects the fraction of true targets required for successful identification in pass 1 DIA queries, in interplay with pass 1 q-value estimation pruning false targets from the empirical library. Whether the 48 residual elastase-attributed precursors are false identifications or non-specific cleavage products of the enzymes present, which the broad elastase rule set would also match, has not been resolved. Under standard two-pass usage, with the empirical library matched to the sample as is the recommended and default mode of operation, cross-enzyme misassignment therefore stays at or below 0.24 % at precursor and peptide level and absent at protein-group level.

While not a precise or complete assessment of error control across multi-enzyme peptide classes, these evaluations, substituting a known-absent peptide class for the unobservable null distribution, suggest that DIA-NN’s error models, although imperfect and not specifically optimized for non-tryptic peptides, nevertheless provide effective error control in the multi-enzyme DIA workflow where minimally a two-pass analysis is performed.

### The mE-DIA Pipeline Runner and mE-DIA Peptide Mapper tools for easy access

Broad adoption of new computational workflows in proteomics often depends not only on methodological performance but also on the availability of tools that make these workflows straightforward to run and interpret. DIA-NN-has demonstrated this convincingly: by combining strong performance with an accessible interface, it has enabled a wide community to employ and interpret DIA-MS workflows with ease and without requiring extensive computational expertise or expensive licensing. In the same spirit, the mE-DIA Pipeline GUI extends the DIA-NN 1.8.1 software suite, the release this pipeline was developed and validated against, with a GUI to orchestrate multi-enzyme analyses that are inaccessible via the native graphical user interface, to lower the practical barriers to multi-enzyme DIA analysis in a broad range of laboratory environments.

The mE-DIA-Pipeline, written in python with full programmatic access, was extended by a react-based GUI and compiled .exe launcher. The GUI enables pipeline set-up, saving and loading of parameter sets, resuming of partial analyses and basic visualization of DIA-NN 1.8.1 results across the enzyme/peptide classes (Fig. S1A). The interface provides visual feedback as each stage completes and presenting the analysis as a connected flow to help users understand the structure of the workflow and monitor progress in real time. To reduce computational overhead and redundant spectral library prediction/generation tasks, the pipeline can detect when spectral library prediction has already been performed in a respective output folder, or, alternatively, pre-computed libraries from earlier analysis iterations can be pinpointed for reuse.

To complement the pipeline runner with a tool for downstream interpretation of multi-enzyme DIA results, we developed the mE-DIA Peptide Mapper, an interactive multi-track peptide mapping viewer with differential discovery and multi-track comparison of condition-specific amino acid observation frequency profiles (Fig. 1C). It takes as input the annotated DIA-NN report generated by the mE-DIA Pipeline together with the corresponding FASTA protein reference database. Its central feature is an interactive peptide map that visualizes peptide observations along individual reference protein backbones, with separate tracks for user-defined conditions to interrogate condition-specific amino acid coverage profiles indicative of alternative isoform usage or shifts in the underlying protein variant composition. Users can toggle the display of semi-specific cleavage peptides and dynamically switch coloring between enzyme, coverage, and track modes to navigate the multi-enzyme data effectively. In addition, a differential discovery module provides a basic scoring scheme to highlight outliers in amino-acid-level coverage difference between a contrast of two conditions, plotted against conventional protein-level fold change or overall abundance; outlier hits can then be inspected directly in the differential peptide map view.

Together, these components form a small but coherent ecosystem that supports the practical dissemination of the mE-DIA-workflow – from raw DIA data, through multi-enzyme analysis, to proteoform-resolved interpretation.

## Discussion

The mE-DIA workflow presented here consolidates multi-enzyme proteomics into a single DIA-MS injection per sample and a single automated analysis run, removing the dependency on per-enzyme DDA-MS measurements, multi-software library construction and manual cross-tool reconciliation that have historically constrained multi-enzyme proteomics to specialized laboratories. By consolidating sample preparation into a pooled C18 cleanup and concentrating all computation within an extended DIA-NN-driven pipeline accessible through a single graphical user interface, the workflow reduces the practical entry barrier and turns multi-enzyme DIA into a routine option alongside conventional tryptic discovery proteomics with minimal additional overhead both experimentally and computationally.

Application to two bacterial proteomes illustrated both the strengths and the present limitations of multiplexed enzyme co-acquisition. In *Streptococcus pyogenes* SF370 and an *Escherichia coli* isolate, the multi-enzyme analyses substantially expanded sequence coverage and the recall of proteoform events relative to the corresponding single-enzyme tryptic workflows, while also incurring losses in per-enzyme peptide identification counts within the multiplex compared to dedicated single-enzyme injections. This trade-off reflects an inherent balance between per-enzyme sensitivity and overall DIA map complexity, with reduced molar loading per enzyme on column, charge competition and ion suppression in the more complex pooled samples likely contributing to attrition in the multiplex. A compelling example of the biological value unlocked by multi-enzyme coverage is the endogenous H69 C-terminal proteoform of the bacterial small ribosomal subunit protein uS19 in *S. pyogenes*, which was invisible in the trypsin-only analysis but resolved through complementary peptide evidence across the TCE multiplex. Given that bacterial ribosomes remain a major antibiotic target in the context of escalating antimicrobial resistance, the ability to pinpoint non-canonical proteoforms and previously hidden molecular surfaces on core translation machinery may open avenues for the discovery of next-generation pathogen-selective therapeutics, while exemplifying how orthogonal enzyme-derived peptide evidence can synergistically expose biologically relevant protein variation that single-enzyme workflows systematically miss.

Operating multi-enzyme DIA acquisitions outside the parameter regime within which DIA-NN’s error models were originally calibrated raises legitimate concerns about FDR validity for non-tryptic peptide classes. The leave-one-out validation framework provides an empirical, if approximate, assessment of cross-enzyme spillover. Under the realistic two-pass default scenario, where the empirical library is rebuilt from the actual sample, raw leakage of analytes from the omitted enzyme class collapsed to zero for three of the four enzyme dropouts and to at most 0.24 % of precursors and peptides, with no protein group, when elastase was omitted, indicating that the workflow does not propagate substantial cross-enzyme error. Uniform peptide-length constraints (7–30 residues) across enzymes and joint scoring within the combined empirical library likely contribute to this robustness, by keeping all peptide classes within the score-distribution regime on which DIA-NN’s q-value estimation is calibrated and effective. Confidence in downstream proteoform calls is further reinforced by gating sequence-divergence events on at least two independent peptide sequence observations (the min_support = 2 default in the proteoform module), which mitigates residual single-peptide false positives, and by discarding supporting precursors attributable to ISD, whose products otherwise pass as non-canonical termini and have been shown to confound proteoform inference (Kim et al., 2013; Schramm et al., 2026).

Benchmarking the library-free mE-DIA workflow against the established DDA-library-based multi-enzyme DIA implementation by Richards et al. on dataset PXD027242 positioned library-free mE-DIA as a competitive alternative. While the experimental DDA library afforded deeper sampling of Glu-C-derived peptides, the library-free workflow identified 20–30 % more precursors, peptides and protein groups within the search space both workflows admit. We cannot rule out that the larger search space of the DDA-based workflow lowered its detection sensitivity, an effect that restricting the comparison to the shared search space after the search does not capture and that we did not quantify. Most striking, however, was the gain in quantitative precision of the library-free route: 74 % more precursors and 47–48 % more protein groups were quantified at ≤ 20 % CV, a gain consistently observed among precursors both workflows quantified (median CV 6.6 % versus 13.8 % in HSC6). Plausibly, though not tested here, it reflects that the pass-1 library is derived directly from the enzyme-multiplexed DIA runs, and thus, by design, captures peptides based on fragment ions, their relative intensities and retention times that are those observed in the enzyme-multiplex DIA data. We surmise, though have not tested, that library-free pass 1 analysis including a first round of interference removal selects fragments for quantification that are less affected by interference from peptides co-isolated in wide-window acquisition. A library built from single-enzyme peptides via narrow-isolation DDA spectra instead can selects fragments by intensity rank in the absence of both, cross-enzyme and co-isolation interference, which may favor selection of fragments that are less suited to quantification in the enzyme-multiplexed wide isolation window DIA data. Importantly, library-free mE-DIA achieves these results without dedicated single-enzyme DDA runs or external library construction, removing a substantial experimental and bioinformatic burden while preserving, and at the protein level extending, quantitative depth.

Several limitations of the present implementation should be acknowledged. The proteoform inference module focuses on a defined subclass of sequence variants, namely non-canonical N- and C-terminal start and stop events along the protein backbone evidenced by two or more independent peptide sequences, within the boundaries of the sequence space annotated in the employed protein sequence database. Other proteoform categories, including chemical modifications, single-amino-acid variants from genetic polymorphism, and internal splice-driven sequence variation, are not directly addressed by the current module and would benefit from dedicated extensions or orthogonal analytical strategies. The events reported here find little corroboration in existing annotation, as merely 6 of the 990 S. pyogenes and 69 of the 544 E. coli termini coincide with an annotated processing boundary in UniProt (all of them initiator-methionine removal or signal-peptide cleavage), whereas this corroboration rate reflects sparse annotation in the database (only 110 of 1,690 and 890 of 4,403 entries carrying any internal processing annotation for *S. pyogenes* and *E. coli*, respectively). This lack of annoted knowledge provides an open field for discovery of protein variants that play biological roles, while potential alternative explanations for non-canonical peptide sequence observations need to be acknowledged and systematically ruled out before selecting candidates for follow-on experimental verification. Terminal peptide evidence cannot inherently distinguish regulated in vivo proteoforms from non-specific proteolytic processing resulting from cell death or routine protein turnover. Consequently, these findings should be viewed as hypotheses for targeted follow-up to separate stable biological signals from degradation products. In addition, the called proteoform events are not fully resolved at the level of the intact molecule, because peptide-level evidence for an N-or C-terminus is grouped per protein but typically only provides locally consecutive information limited by the maximal peptide length sampled. It therefore remains unknown to what extent multiple proteoform events co-occur on the same molecule and what the underlying full-length proteoform backbones are. Top-down and middle-down proteomics workflows are uniquely positioned to dissect such intramolecular co-occurrence, and we anticipate a fruitful synergy space in which multi-enzyme bottom-up peptide maps are referenced against intact protein backbones and middle- or top-down-derived proteoform inventories to recover combinatorial proteoform identities at scale. Computationally, the multi-enzyme, multi-pass workflow is also considerably more demanding than conventional fully-specific single-enzyme DIA analysis. In particular, the semi-specific extension step (passes 3 and 4) expands the searched peptide space substantially and, for studies covering large numbers of confidently identified protein groups, poses a challenge for classical consumer-grade hardware, in practice requiring strong workstation configurations or HPC execution to complete in reasonable time.

Two asymmetries in the acquisition itself are worth addressing in future implementations, and both are visible in the per-enzyme peptide properties. Although the digests were combined in equal proportions (by input protein amount and volume here, 1:1:1 by weight in the human data), tryptic peptides accounted for 52.9 % of the total normalized precursor signal in the four-enzyme *E. coli* multiplex, 68.6 % in the three-enzyme *S. pyogenes* multiplex and 75.8 % in the three-enzyme human runs (Fig. S2C), so the effective sampling depth per enzyme is not equal and pooling digests in proportions calibrated on measured signal share, rather than on input amount, could help distribute sampling depth more equally across different peptide classes. Second, retention-time prediction is less accurate for non-tryptic peptides, illustrating how predictors trained on tryptic peptide sets give non-tryptic peptide classes a systematically weaker prior, outlining opportunities for improved non-tryptic peptide detection by retraining or fine-tuning predictors on multi-enzyme data would benefit identification directly (Fig. S2E).

These results establish library-free mE-DIA as a self-contained, scalable workflow well suited for differential quantitative discovery across diverse enzyme combinations. The present implementation is deliberately anchored in DIA-NN 1.8.1, the release mE-DIA was developed and validated against, which remains freely available for both, academic and also commercial use. Future efforts may ensure seamless extendability to more recent DIA-NN releases that offer better predicted libraries and support larger search spaces and improved FDR control. Extension to newer engines with expanded PTM coverage, integration with proteogenomic reference databases derived from RNA-seq or ribosome profiling data, and benchmarking on next-generation high-throughput instrumentation (timsTOF HT, Orbitrap Astral) are natural next steps that should further broaden the proteoform landscape accessible from a single bottom-up multi-enzyme DIA injection.

## Methods

### Bacterial strains and culture conditions

*Streptococcus pyogenes* strain SF370 was grown in Todd-Hewitt broth supplemented with 0.5% (w/v) yeast extract (THY-media). The bacteria were grown at 37 °C, 5% CO2 until mid-logarithmic phase (OD620 nm 0.4–0.5), and harvested by centrifugation (3500g, 5 min, RT), prior to cell lysis and sample preparation for mass spectrometry. The cells were lysed in 8M urea-100 mM ammonium bicarbonate, with subsequent sonication using a Bioruptor Plus (Diagenode) at 4 °C.

*Escherichia coli* strain O18K1 was grown in Luria-Bertani (LB) broth. The bacteria were grown at 37 °C, 200 rpm, until mid-logarithmic phase (OD600 nm 0.5). For protein extraction, pellets were centrifuged at 13,000 xg, the supernatant removed and samples air-dried before re-suspension in 6 M guanidine and 50 mM Tris-HCl lysis buffer, prior to sonication using the Bioruptor Plus at 4 °C.

### Sample Preparation for Mass Spectrometry

*E. coli* and *S. pyogenes* cell pellets were lysed by bead-beating (3 × 2 min, with cooling at 4 °C between cycles) in a buffer containing 8 M urea and 100 mM ammonium bicarbonate, followed by denaturation and disulfide bond reduction with 5 mM Tris(2-carboxyethyl)phosphine hydrochloride (TCEP) for 60 min at 37 °C, shaking at 800 rpm. Free cysteines were alkylated with 10 mM iodoacetamide for 30 min in the dark at room temperature. Samples were then diluted to a urea concentration <1.5 M with 100 mM ammonium bicarbonate prior to protein concentration determination by OD280 and sample splitting for per-enzyme digests at a Enzyme:Substrate ratio of ≈1:50 at 37 °C, shaking at 800 rpm overnight. Multi-enzyme digestion was performed in parallel: aliquots of the sample were digested separately with trypsin (V5111), chymotrypsin (V1062), Glu-C (V1651) and elastase (V1891), all enzymes from Promega, under standard conditions. Following digestion, reactions were quenched by adjusting the pH to ≤ 2 by addition of formic acid before any digests were combined, so that no protease could act on the peptides of another digest. To support the ensuing leave-one-out experimental design to explore error propagation, quenched individual enzyme digests were not pooled but desalted individually using C18 solid-phase extraction and combined into either one or leaving-one-out pooled samples. As a side note for production mode experiments, pooling before C-18 clean-up minimizes consumable cost and hands-on preparation time compared to per-enzyme cleanup and concatenation.

### LC-MS/MS Acquisition

Samples were analyzed on an Thermo Fisher Orbitrap HF-X mass spectrometer coupled with reversed-phase liquid chromatography on a EASY-nLC 1200, using a EasySpray column with integrated emitter, temperature-controlled at 45 °C (Thermo Fisher ES902), and separation gradients from 5 %-40 % Buffer B (0.1 % formic acid in 80 % Acetonitrile) in Buffer A (0.1 % formic acid) at 350 nl/min flow rate over 60 min (E.coli, dataset 1) or 120 min (S. pyogenes, dataset 2). Scans at resolutions of 120,000 (MS1) covered a precursor range of 350-1650 m/z coupled to 26 variably sized DIA windows for binned MS2 acquisitions (350–383, 382–408, 407–429, 428–448, 447–467, 466–484, 483–503, 502–521, 520–539, 538–557, 556–575, 574–594, 593–614, 613–634, 633–656, 655–678, 677–701, 700–726, 725–756, 755–787, 786–823, 822–862, 861–914, 913–979, 978–1077, and 1076–1650 m/z). The 60-min gradient runs employed a MS2 resolution of 30,000, whereas the 120-min runs employed R = 45,000. For both gradients, MS1 scans were acquired at a resolution of 120,000 across 350–1650 m/z, with an AGC target of 3e6, a maximum injection time of 100 ms, and profile-mode data acquisition. MS2 (DIA) scans used an AGC target of 3e6 and HCD fragmentation at a fixed normalized collision energy (NCE) of 27, with a fixed first mass of 140 m/z, a loop/multiplexing (MSX) count of 1, and profile-mode acquisition, across the same 26 variable-width windows spanning 350–1650 m/z with 1 m/z overlap. In the 60-min runs, MS2 scans were acquired at R=30,000 with a maximum injection time of 50 ms, giving a maximal duty cycle of ≈2.3 s, whereas 120-min runs afforded MS2 scans at R=45,000 with a maximum injection time of 100 ms, giving a maximal duty cycle of ≈3.6 s.

### Multi-Enzyme Library-free DIA Search and Candidate Proteoform Inference

Peptide identification was performed using DIA-NN version 1.8.1, orchestrated by the mE-DIA pipeline GUI. In Pass 1, an independent library search was conducted for each enzyme with --cut parameter set to the enzyme-specific cleavage rule (Table 1), --missed-cleavages 1 (3 for elastase), --fixed-mod UniMod:4,57.021464,C (carbamidomethylation of cysteine), --met-excision, --min-pep-len 7, --max-pep-len 30, --out-lib, --full-profiling, and --smart-profiling. The peptide length window of 7–30 amino acids was applied uniformly across all enzymes, and no variable modifications were searched. Protein sequence databases were UniProt reference proteomes with one sequence per gene, retrieved 2026-01 (filename tag Date202601): *S. pyogenes* SF370 (UP000000750, 1,690 entries), *E. coli* K-12 (UP000000625, 4,403 entries) and human (UP000005640, 20,659 entries).

**Table 1:** Enzyme cleavage rules implemented in meDIA pipeline.

| Code | Enzyme | --cut rule | Sense | Missed cleavages (default setting) |
|---|---|---|---|---|
| T | Trypsin | K*,R*,!*P | C-terminal | 1 |
| T2 | Trypsin/P | K*,R* | C-terminal | 1 |
| C | Chymotrypsin | F*,W*,Y*,L*,!*P | C-terminal | 1 |
| G | Glu-C | D*,E*,!*P | C-terminal | 1 |
| EX / E-slim | Elastase, extended (used here) / restricted | A*,V*,S*,G*,L*,I* / A*,V* | C-terminal | 3 |
| L | Lys-C | K*,!*P | C-terminal | 1 |
| LN | LysN | *K | N-terminal | 1 |
| R | ArgC | R*,!*P | C-terminal | 1 |
| A | Asp-N | *D | N-terminal | 1 |
| M | CNBr | M* | C-terminal | 1 |
| F | PepsinA | F*,L*,!*P | C-terminal | 1 |

The enzyme-specific empirical spectral libraries were merged into a composite library by concatenating their contents, deduplicating entries by transition_group_id, and annotating each entry with an Enzyme column. Deduplication by transition_group_id keeps one entry per precursor, so a peptide that more than one enzyme can generate is represented once and carries a single enzyme attribution; the Enzyme column therefore records which enzyme-specific library contributed the entry, not that only that enzyme can produce the peptide, and per-enzyme stratifications inherit that attribution. In Pass 2, the composite library was used for a final DIA-NN search with --cut "" (disabled, as library entries are already enzyme-annotated), --matrices, and --matrix-qvalue 0.01, generating quantification matrices at precursor, peptide, and protein group levels at 1% FDR.

For semi-specific extension (Passes 3 and 4), analysis was constrained to protein sequences identified with Global.PG.Q.Value ≤ 0.01 in Pass 2, followed by in silico library generation via DIA-NN’s built-in predictor for this subset protein sequences, employing semi-specific cleavage rules. These semi-specific in silico generated libraries were then queried against the DIA data per each enzyme and peptide class to generate the semi-specific empirical libraries. Pass 4 merged the semi-specific libraries across enzymes with the Pass 2 composite library (fully specific peptides across enzymes) and re-searched the raw DIA data, producing a final report_annotated.tsv extended with columns Enzyme, Specificity, Sequence.Start and Sequence.Stop for simplified downstream multi-enzyme peptide mapping analyses. Sequence-divergence events were called by the pipeline’s proteoform module, which requires at least two distinct peptide sequences sharing a free terminus at the same residue (min_support = 2 default); one event is one terminus at one residue, so a start and a stop at the same position count separately, and the number of proteases contributing to an event was taken from the Enzyme annotation of its supporting peptides. Supporting precursors attributable to ISD were then discarded. A precursor was flagged when an identified longer peptide sharing its event-distal terminus (the same C-terminus for N-terminal events, the same N-terminus for C-terminal events) had a precursor with median apex retention time within 0.05 min and a higher median precursor quantity. A peptide sequence supported an event only through unflagged precursors, and events left with fewer than two supporting sequences were removed; flagged precursors are kept in the proteoform matrix with their parent peptide (columns isd_flag and isd_parent) for inspection. The proteoform step of every analysis reported here was run with mE-DIA v1.1.0.

### Candidate Proteoform Event Review Against Public Reference Database

Detected termini were compared against UniProt processing annotation for the same proteome (initiator methionine, signal and transit peptide, propeptide, mature chain and peptide features; retrieved 2026-09-18). A corroboration was counted when a C-terminal event coincided with the last residue of such a feature, or an N-terminal event with its first residue or with the residue following an annotated cleavage; boundaries of chain features spanning the full entry length were treated as canonical protein termini and not counted as processing evidence.

### Reanalysis of the Richards et al. Multi-Enzyme DIA Dataset (PXD027242)

The six multiplex DIA runs of the HCD acquisition series (Y20210522-29, -31, -33 for HSC6 and Y20210522-35, -37, -39 for SCC25; three replicates per cell line; trypsin / Glu-C / Asp-N) were obtained from PRIDE (PXD027242) and searched with the library-free mE-DIA workflow described above, against the human UniProt reference proteome with one sequence per gene (UP000005640, 20,659 entries, 1 sequence per gene downloaded 01/2026) and with the pass-1 and pass-2 parameters described above except that up to two missed cleavages were allowed, to approximate the settings of the DDA-based workflow. The semi-specific extension was not applied to this dataset, so every comparison reported here is between fully specific searches, as reported by (Richards et al., 2022). The DDA-based side of the comparison was not re-searched: it is the deposited DIA-NN report and quantitative matrices for the same six raw files (DLS24_Mix_HCD_report_DIANN.tsv, DLS24_Mix_HCD_report.pr_matrixL.tsv, DLS24_Mix_HCD_report.pg_matrix.tsv) together with the deposited merged DDA spectral library (DLS24_merge_file_Library_HCD.csv), all part of PXD027242. Both sides were produced by DIA-NN from identical raw data. They differ in three respects: the spectral library queried (the authors’ DDA library versus the mE-DIA empirical pass-1 library derived from the DIA data itself); the settings that define the search space (enzyme rules, missed cleavages, peptide length, precursor charge and variable modifications); and the DIA-NN version and quantification settings (Richards et al. used DIA-NN 1.7.16). We tested the last directly: re-searching the six raw files with the deposited DDA library under DIA-NN 1.8.1 and the settings used here reproduced the deposited identifications to within 1.4 % at precursor and peptide level and 0.4 % at protein-group level. The search space of a workflow comprises all peptides its settings admit from its FASTA. For library-free mE-DIA it was enumerated from the logged pass-1 settings (trypsin after K or R and Glu-C after D or E, neither before P; Asp-N before D; up to two missed cleavages; 7–30 residues; fixed carbamidomethylation; initiator methionine excision; no variable modifications; precursor charge 1–4 and m/z 300–1,800), which reproduced the numbers of precursors DIA-NN generated per enzyme to within 0.02 %. No search parameters were deposited for the DDA-based workflow, so its search space was inferred from the content of the deposited library: Swiss-Prot human canonical sequences (approximated by the Swiss-Prot entries of the FASTA used here), Glu-C after E only, up to three missed cleavages, 7–52 residues, fixed carbamidomethylation, variable methionine oxidation and protein N-terminal acetylation (up to five per peptide) and initiator methionine excision. The shared search space contains the unmodified precursors whose sequence lies in both search spaces, at charge 1–4 and m/z 300–1,800; DDA-library precursors with variable modifications or charge above 4 fall outside it by definition. All library, identification and precision comparisons were restricted to it, matching precursors by stripped sequence and charge. Identifications count analytes detected in all three replicates of a cell line at 1 % FDR, unioned across the two cell lines; a protein group was counted when at least one of its precursors within the shared search space met this rule, at DIA-NN’s protein-group q-value of 1 %. Because protein-group quantities cannot be restricted to a subset of precursors after the search, they were recomputed for both workflows from the restricted precursor matrices with the MaxLFQ algorithm (median pairwise log-ratios solved by least squares, (Cox et al., 2014)). Applied to the unrestricted matrices, this re-quantification reproduced DIA-NN’s protein-group profiles (within-protein Pearson r of log2 quantities 0.993 and 0.997 for the library-free and DDA-based results) and the number of HSC6 protein groups at ≤ 20 % CV to within 1 %. Coefficients of variation were computed across the three replicates of a cell line from each workflow’s own precursor and protein-group matrices (≥ 2 non-zero values). Library contents were compared per enzyme channel on stripped and modified peptide sequences.

### FDR Estimation and Leave-One-Out Validation

To empirically assess cross-enzyme spillover, each enzyme was systematically omitted from the sample (conditions dT, dC, dG, dE) while retaining its entries in the pass 1 TCGE composite library, then used to search each dX sample (probing leakage in “pass 2 only”), and a second mode (“pass 1 + pass 2”), in which the full pipeline of pass 1 search and pass 2 search were (re-)run on each dX sample to generate a sample-matched empirical library. Raw leakage was reported as the fraction of total IDs attributable to the omitted enzyme, and library leakage as the fraction of omitted-enzyme entries in the library passing identification thresholds. Protein groups (semicolon-delimited Protein.Ids) and respective ambiguous peptide level evidence were excluded from LooV counting.

### Enzyme Cleavage Rules

DIA-NN’s --cut parameter was configured for each enzyme according to standard protease/cleavage specificities (Table 1). The syntax encodes cut positions with an asterisk and blocking neighbors with an exclamation mark prefix. Data from five primary enzymes were used in this work; seven additional enzymes are supported in the mE-DIA Pipeline registry. For elastase, the extended rule set A, V, S, G, L, I (mE-DIA code EX) was used with up to three missed cleavages; throughout the text and figures, ‘elastase’ refers to this rule set. The pipeline additionally provides the restricted A, V rule (E-slim).

### Software and Algorithms

#### mE-DIA Pipeline

The mE-DIA Pipeline is a Python (≥ 3.10) wrapper around DIA-NN (v1.8.1), which performs all spectral library prediction and search passes (See above and Results section). Supporting modules handle the enzyme cleavage-rule registry, library merging, and annotation of the final report with each precursor’s enzyme of origin, its specificity (recomputed from the protein sequence) and its sequence position. A final module removes in-source decay peptides and collects proteoform-differentiating events and supporting evidence. Data handling uses pandas, and plots use matplotlib. The graphical user interface pairs a FastAPI/Uvicorn backend with a React/TypeScript front end styled with Tailwind CSS and built with Vite. The interface covers (i) configuration of DIA-NN executable and pipeline input raw files, FASTA database, enzyme and other search parameters and pipeline output folder (ii) monitoring pipeline progress pass by pass with a live view of the pipeline’s console output and (iii) results view of per-run precursor, peptide and protein-group identification counts.

#### mE-DIA Peptide Mapper

The Peptide Mapper is a standalone, single-file HTML/JavaScript application, built with React, Tailwind CSS and Lucide icons. The pipeline writes it into each result folder, and it is opened separately in a web browser. It reads the annotated report and the FASTA database used for the search. Optionally, runs can be assigned to user-defined groups. The PeptideMap module draws all peptides of a selected protein along its sequence, as one track for all data, per run or per group. Peptides are coloured by enzyme, by uniqueness or by track, and semi-specific peptides are highlighted. A residue-level observation-frequency profile is shown for each analysis group and feeds the Differential Discovery module. It ranks proteins by a discordance score S, the summed absolute difference between two groups’ residue-level observation frequencies, i.e. the number of peptide observations covering each residue summed over a group’s runs and divided by the number of runs in that group. S is plotted against either the log₂ fold change of mean protein-group quantity between the two groups or the mean protein-group quantity across both groups. Only proteins with peptide evidence in both groups are scored. S is a ranking aid, not a statistical test, and is not normalised for protein length.

### Software and Data Availability

The mE-DIA Pipeline requires a functional DIA-NN installation and is made available both as a command-line tool (https://github.com/heuselm/mE-DIA/blob/v1.1.0/bin/run_pipeline.py) and as a graphical user interface via a self-contained Windows executable bundling the server, the user interface, a Python 3.14 runtime and all Python dependencies (https://github.com/heuselm/mE-DIA/releases/tag/v1.1.0). The DIA Peptide Mapper is a lightweight, standalone HTML/JavaScript application which the pipeline writes into each result folder and which runs without installation in the user’s web browser. For separate use, e.g. to generate interactive peptide maps from conventional DIA-NN result files, the viewer is available at https://github.com/heuselm/mE-DIA/blob/v1.1.0/bin/viewer/viewer.html. Raw mass spectrometry data including sdrf annotation (Dai et al., 2021) and processed data archives have been deposited to ProteomeXchange via the PRIDE partner repository (https://www.ebi.ac.uk/pride/archive/projects/PXD085278/ ; will be made available to reviewers & upon acceptance). Richards et al. raw data and result files used here are available from the original deposition (PXD027242, (Richards et al., 2022)).

## Author contributions

The manuscript was written through contributions of all authors. M.H. conceived and implemented the mE-DIA workflow; performed analysis and implemented the mE-DIA Pipeline and Peptide Mapper software. L.H. performed the MS sample preparation and MS analysis. C.K. co-developed software, performed MS maintenance and analysis. E.Å. performed MS maintenance and analysis and gave critical input. J.M. supervised the study. All authors have given approval to the final version of the manuscript.

## Acknowledgements

We would like to acknowledge fruitful discussions on multi-enzyme de novo sequencing and computational frameworks with Carlos Gueto Tettay, Aaron Scott and Lars Malmström. This study was supported by funding from The Knut and Alice Wallenberg Foundation, the Medical Faculty at Lund University and Region Skåne, as well as the Strong Research Environment MultiPark at Lund University.

