## Supplemental Information for "Multi-Enzyme Data-Independent Acquisition Mass Spectrometry for Proteoform-Aware Single-shot Proteomics"

### Supplementary Figures

This document contains Supplementary Figures S1–S5 supporting the main manuscript. The analyses, enzyme abbreviations and the ISD filter are described in the Methods of the main manuscript.

Figure S1: mE-DIA Pipeline interface and precursor-level evidence for P66496_H69C

**A** **mE-DIA Pipeline user interface**

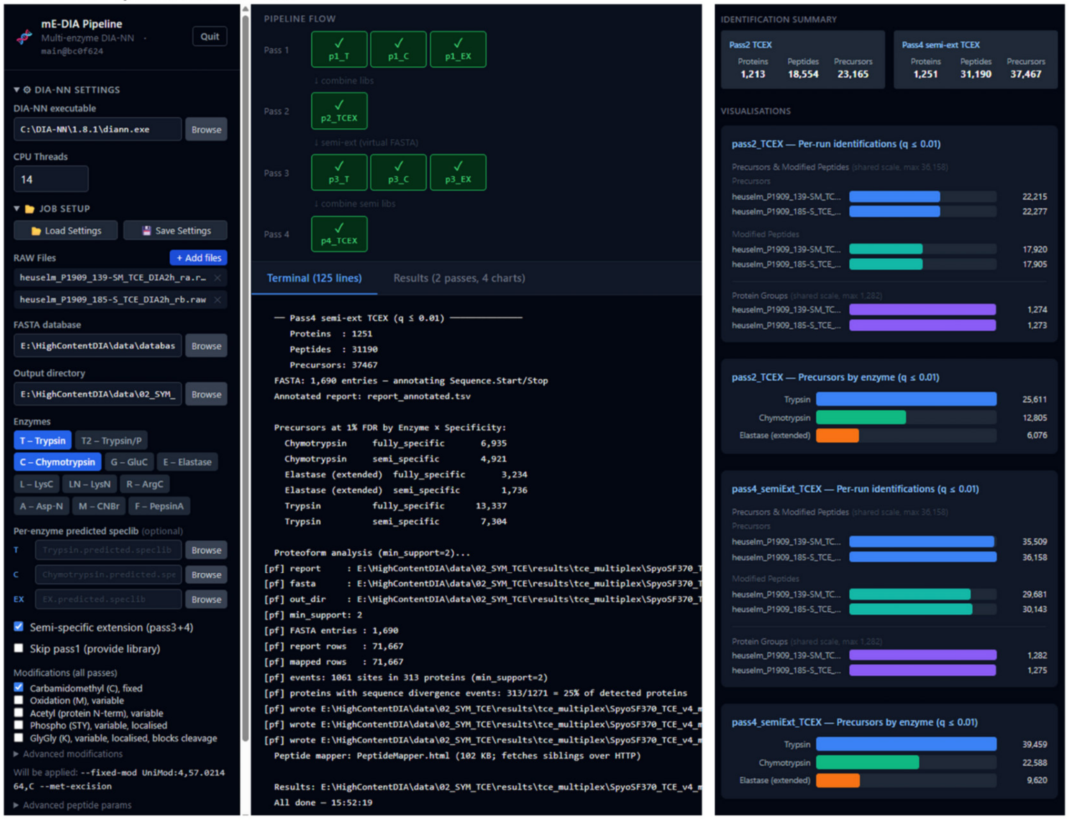


**B** **Precursor-level evidence, uS19 H69 C-terminus**

***S. pyogenes* SF370, P66496_H69C — 30S ribosomal protein uS19, C-terminus at residue 69**
**5 distinct sequences, 8 precursors, 3 proteases**

| Peptide | Charge | Protease | Residues | q-value |
|---|---|---|---|---|
| DGRKHVPVYIQEDMVGH | 2 | Chymotrypsin | 53–69 | 5.7 E-4 |
| DGRKHVPVYIQEDMVGH | 3 | Chymotrypsin | 53–69 | 8.0 E-5 |
| DGRKHVPVYIQEDMVGH | 4 | Chymotrypsin | 53–69 | 3.0 E-4 |
| IQEDMVGH | 2 | Chymotrypsin | 62–69 | 5.7 E-4 |
| YIQEDMVGH | 2 | Elastase | 61–69 | 4.0 E-4 |
| HVPVYIQEDMVGH | 3 | Trypsin | 57–69 | 2.9 E-4 |
| KHVPVYIQEDMVGH | 3 | Trypsin | 56–69 | 1.7 E-4 |
| KHVPVYIQEDMVGH | 4 | Trypsin | 56–69 | 2.9 E-3 |

Figure S1: **The mE-DIA Pipeline graphical user interface, and precursor-level evidence for the uS19 H69 C-terminal proteoform. (A)** The mE-DIA Pipeline graphical user interface after a completed four-pass analysis of the *S. pyogenes* SF370 TCE multiplex: DIA-NN settings, job setup, enzyme selection and modifications (left), the pipeline flow per pass with the run log (centre), and per-run identification summaries (right). The proteoform line of the run log was recorded before the ISD filter was introduced and reports the unfiltered event count. **(B)** Precursors supporting the P66496_H69C call (30S ribosomal protein uS19, C-terminus at residue 69) in the TCE multiplex, pass 4, with charge, protease, residue range and experiment-wide precursor q-value; none of the eight precursors is flagged as an ISD product.

Figure S2: Peptide properties and chromatography metrics (1 % FDR)

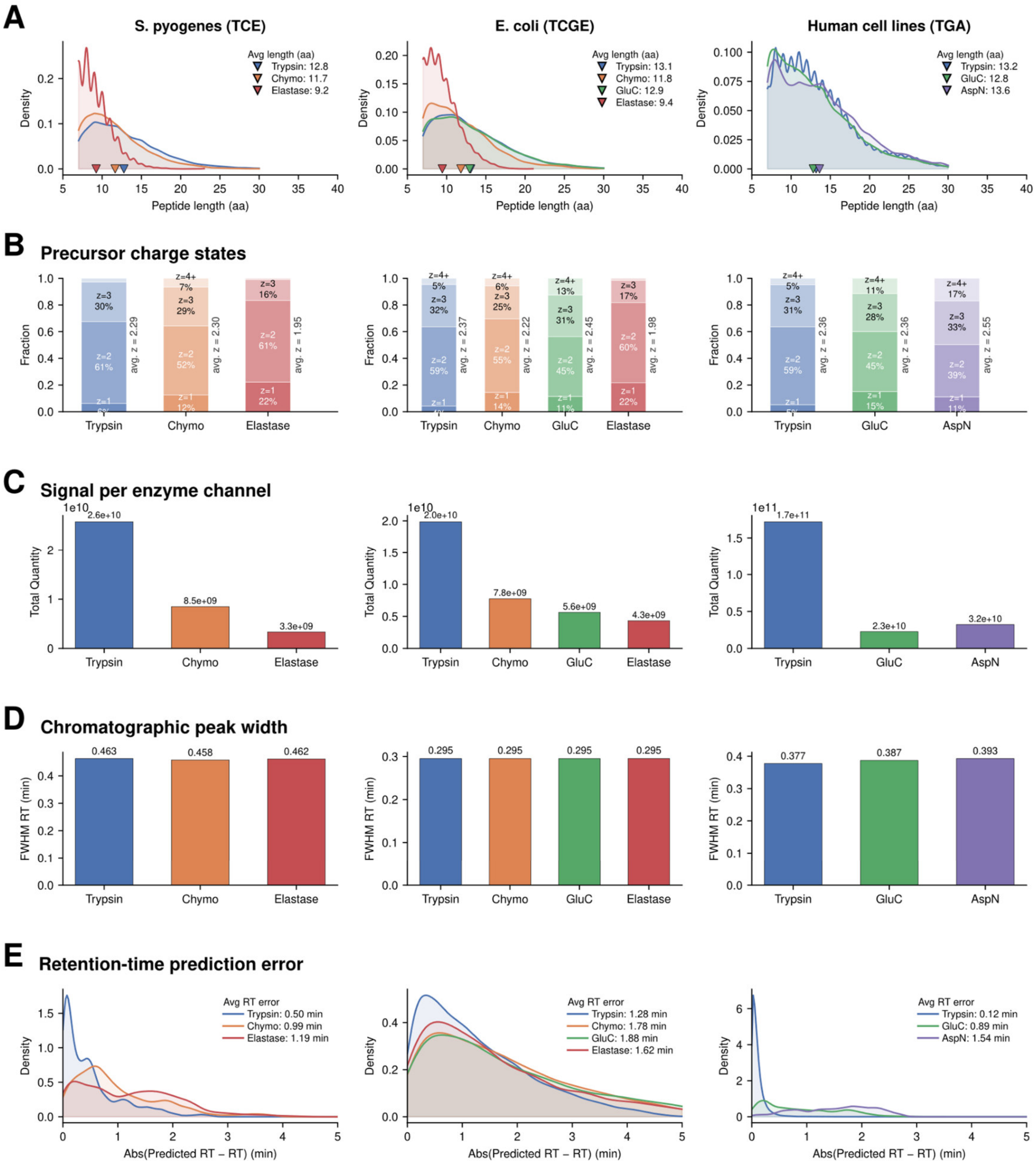


Figure S2: **Peptide and chromatographic properties per enzyme channel.** Columns: *S. pyogenes* SF370 (TCE), *E. coli* O18:K1 (TCGE) and the human cell lines of PXD027242 (TGA); precursors at 1 % FDR. **(A)** Peptide length distribution, with the mean length per enzyme. **(B)** Precursor charge-state distribution, with the mean charge. **(C)** Total precursor quantity per enzyme channel; digests mixed 1:1:1(:1) by volume do not contribute equally to the signal. **(D)** Median chromatographic peak width (FWHM), indistinguishable between enzyme channels within a dataset. **(E)** Absolute deviation between observed and predicted retention time, larger for the non-tryptic peptide classes.

Figure S3: TCE-Recall of analytes in TCE multiplex mE-DIA relative to separate per-enzyme T/C/E DIA (*S. pyogenes*)

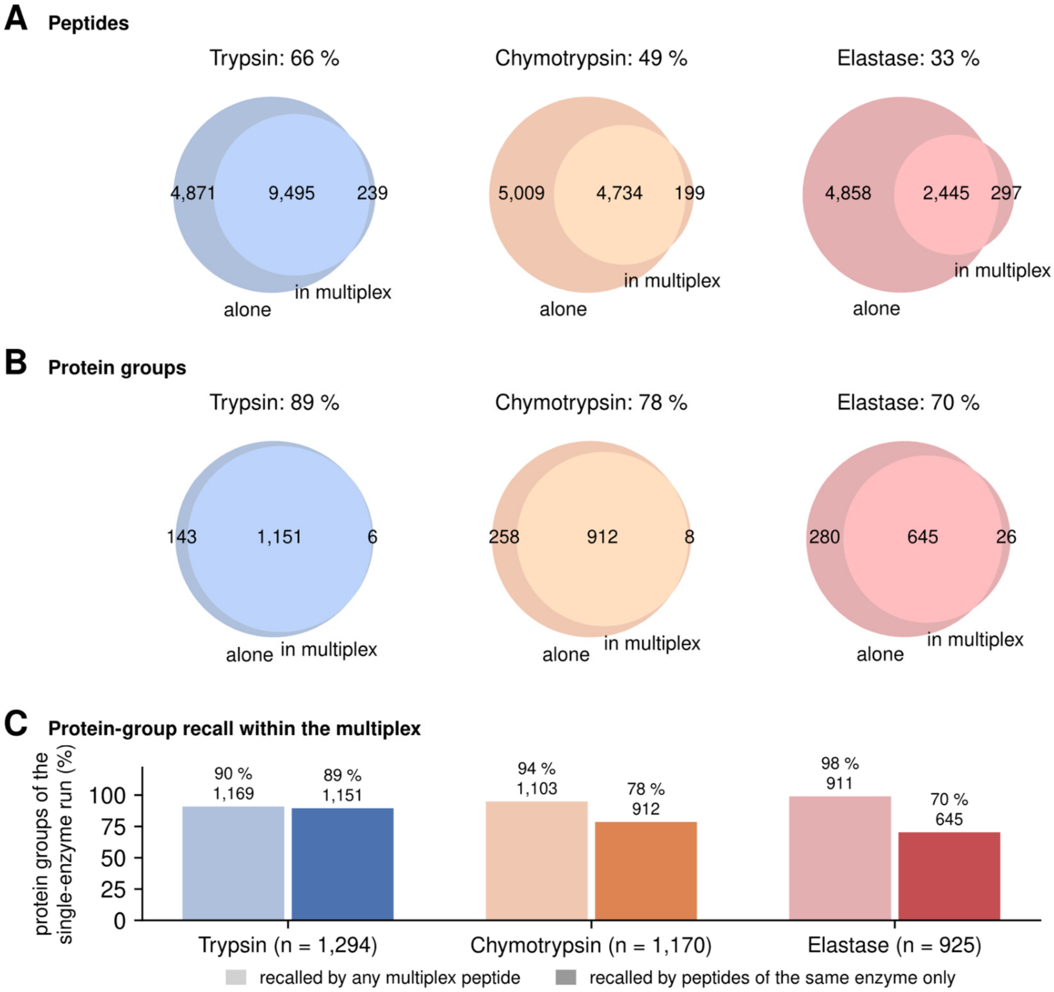


Figure S3: **Per-enzyme recall within the *S. pyogenes* SF370 TCE multiplex, each single-enzyme analysis against the same enzyme's channel of the multiplex.** Fully specific pass 2 on both sides; identifications in both replicates at 1 % FDR. A multiplex precursor belongs to the channel of the enzyme that produced it in the merged pass-1 library. **(A, B)** Two-set Venn diagrams of each single-enzyme analysis (alone) against the peptides of that enzyme within the multiplex (in multiplex), at peptide **(A)** and protein-group **(B)** level; protein groups in **(B)** are those supported by peptides of the respective channel. Diagram titles give the share of the single-enzyme identifications recalled in the multiplex. **(C)** Protein-group recall within the multiplex when any multiplex peptide may support a protein group (light bars) and when only peptides of the same enzyme may (dark bars); n = protein groups of the single-enzyme analysis.

Figure S4: Semi-specific extension gains, quantitative reproducibility in multiplex and event detection rate analysis vs. per-enzyme DIA combined data

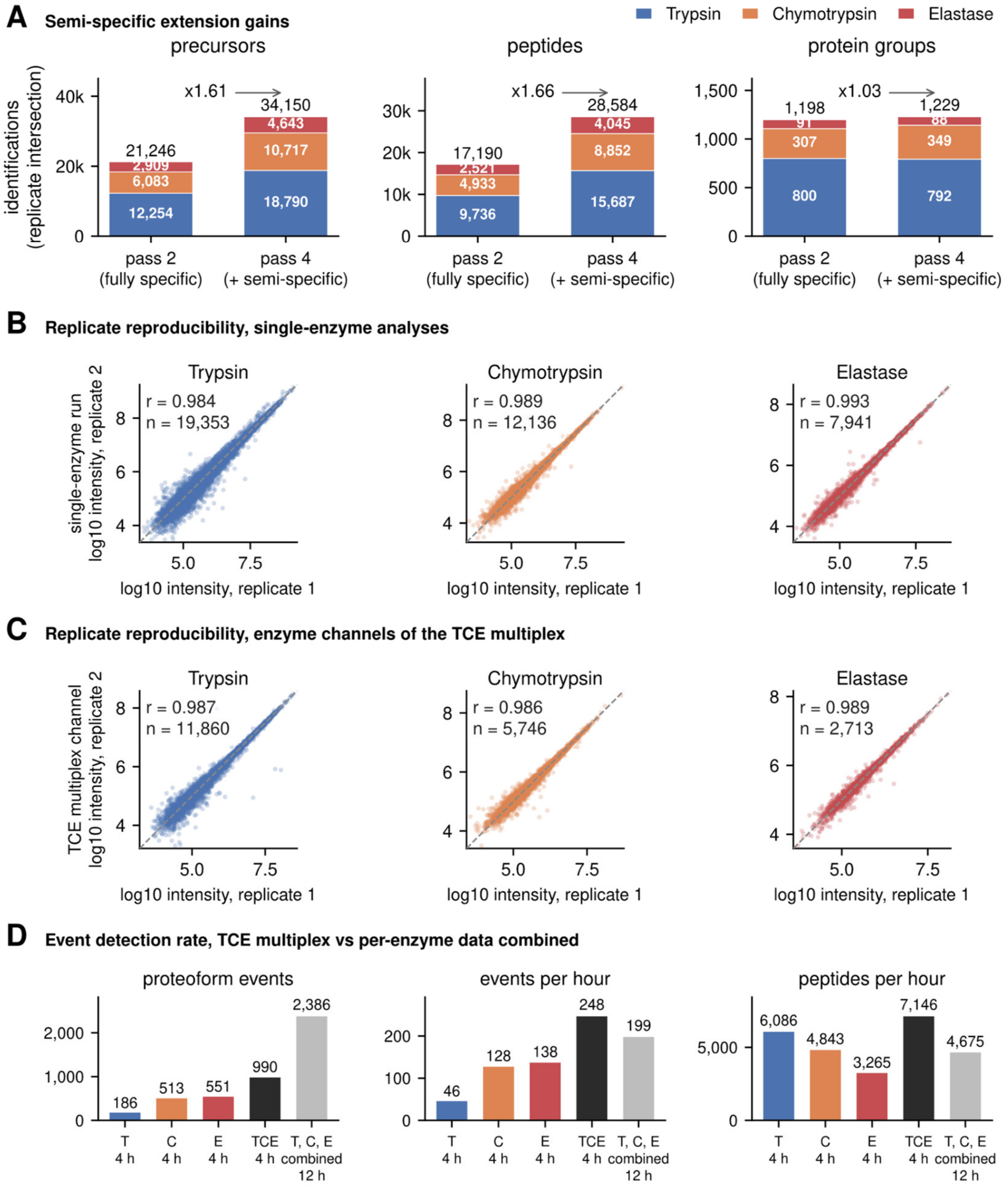


Figure S4: **Semi-specific gain, replicate reproducibility, and comparison with a pseudo-pool for the *S. pyogenes* SF370 TCE multiplex. (A)** Identifications in the fully specific pass 2 and the semi-specific extended pass 4, stacked by enzyme channel, at precursor, peptide and protein-group level (replicate intersection at 1 % FDR), with the pass-4 over pass-2 ratio annotated. **(B, C)** Precursor-level replicate correlation (log10 intensity, replicate 1 against replicate 2) in the individual single-enzyme analyses **(B)** and for the peptides of the same enzyme within the multiplex **(C)**, with Pearson r and the number of precursors. **(D)** The physical TCE multiplex compared with the data of the three single-enzyme analyses combined (T, C, E combined): identified separately, then passed together through the same proteoform inference including the ISD filter: candidate proteoform events, events per hour of acquisition and peptides per hour. The same volume (6 µL) was injected for every run, so each digest is loaded at one third in the multiplex. Acquisition time is two replicates × 2 h per analysis; peptides are those identified in both replicates (pass 4).

Figure S5: Proteoform inference across analyses, enzyme channels and multiplexes, and the ISD filter

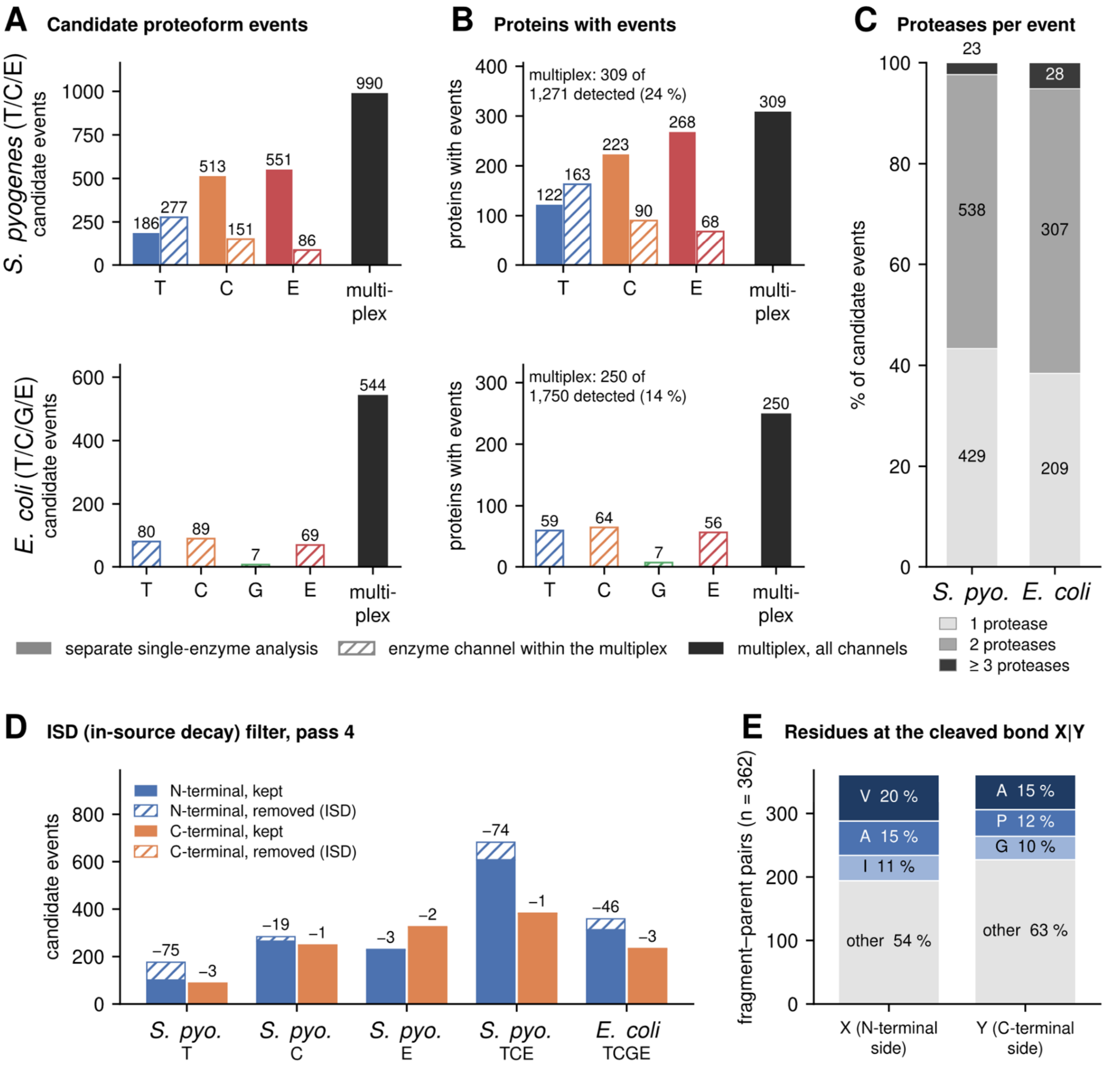


Figure S5: **Proteoform inference across single-enzyme analyses, enzyme channels and multiplexes, and the effect of the ISD filter.** Pass 4. **(A)** Candidate proteoform events in *S. pyogenes* SF370 (top) and *E. coli* O18:K1 (bottom): separate single-enzyme analyses (solid bars), enzyme channels within the multiplex (hatched bars; the multiplex re-scored with that enzyme's peptides only, under the same two-sequence rule, not separate experiments) and the multiplex with all channels (black). No single-enzyme analyses were acquired for *E. coli*. **(B)** Proteins with at least one event, with the multiplex count relative to the detected proteins. **(C)** Number of proteases contributing supporting peptides per multiplex event (1, 2, ≥ 3). **(D)** Candidate events kept and removed by the ISD filter per analysis and terminus, with the number removed above each bar. **(E)** Residues on either side of the bond cleaved in the source, X|Y, over all 362 unique fragment–parent pairs (N- and C-terminal events, both organisms pooled): the three most frequent residues per side, the remainder as other; labels give the share of pairs.
